# An adaptable plasmid scaffold for CRISPR-based endogenous tagging

**DOI:** 10.1101/2023.11.01.565029

**Authors:** Reuben Philip, Amit Sharma, Laura Matellan, Anna C. Erpf, Wen-Hsin Hsu, Johnny M. Tkach, Haley D. M. Wyatt, Laurence Pelletier

**Affiliations:** Lunenfeld-Tanenbaum Research Institute, Mount Sinai Hospital, Toronto, Ontario, M5G 1X5, Canada; Department of Molecular Genetics, University of Toronto, Toronto, Ontario, M5S 3E1, Canada; Department of Biochemistry, University of Toronto, Toronto, Ontario, M5S 1A8, Canada

**Keywords:** Gene-editing, CRISPR, Endogenous Tagging

## Abstract

Endogenous tagging makes it possible to study a protein’s localization, dynamics, and function within its native regulatory context. This is typically accomplished via CRISPR, which involves inserting a sequence encoding a functional tag into the reading frame of a gene. However, this process is often inefficient. Here, we introduce the “quickTAG,” or qTAG system, a versatile collection of optimized repair cassettes designed to make CRISPR-mediated tagging more accessible. By including a desired tag sequence linked to a selectable marker in the cassette, integrations can be quickly isolated post-editing. The core sequence scaffold within these constructs incorporates several key features that enhance flexibility and ease of use, such as: specific cassette designs for N- and C-terminus tagging; standardized cloning sequences to simplify the incorporation of homology arms for HDR or MMEJ-based repairs; restriction sites next to each genetic element within the cassette for easy modification of tags and selectable markers; and the inclusion of lox sites flanking the selectable marker to allow for marker gene removal following integration. We showcase the versatility of these cassettes with a diverse range of tags, demonstrating their applications in fluorescence imaging, proximity-dependent biotinylation, epitope tagging, and targeted protein degradation. The adaptability of this scaffold is also exhibited by incorporating novel tags such as mStayGold, which offer enhanced brightness and photostability, reconciling prolonged live-cell imaging of proteins at their endogenous levels. Finally, by leveraging the restriction sites, entirely distinct cassette structures and editing schemes were developed. These enabled scenarios that included conditional expression tagging, selectable knockout tagging, and safe-harbor expression. Our existing and forthcoming collection of plasmids will be accessible through Addgene. It includes ready-to-use constructs targeting common subcellular marker genes, as well as an assortment of tagging cassettes for the tagging of genes of interest. The qTAG system offers an accessible framework to streamline endogenous tagging and will serve as an open resource for researchers to adapt and tailor for their own experiments.

## Introduction

Gene overexpression has long been a crucial tool in molecular biology, offering insight into gene function and how genes contribute to disease. Nevertheless, this approach has its constraints. Excessive protein abundance can result in mislocalization, causing interactions to deviate from their native state and ultimately manifest in ectopic phenotypes^1–5^. This emphasizes the importance of examining proteins at their native expression levels where a more accurate comprehension of a protein’s localization, dynamics, and interactions can be studied. Since its discovery in 2012, the advent of Clustered Regularly Interspaced Short Palindromic Repeats (CRISPR) alongside its programmable Cas endonucleases has sparked a transformative revolution in molecular genetics^6^. Using a single guide RNA (sgRNA), this technology enables the editing of DNA by directing the Cas endonucleases to a specific genomic locus and allowing for targeted edits to be made. This is especially relevant in the context of endogenous tagging, where the insertion of large transgenes encoding functional tags into the reading frame of a gene of interest (GOI) facilitates the study of proteins, without the potential consequences of overexpression.

Cas endonucleases, such as Cas9, exhibit a high level of precision in identifying DNA sequences to cleave. When guided by the sgRNA, Cas9 locates a DNA sequence matching the targeting sequence adjacent to a protospacer adjacent motif (PAM). This prompts Cas9 to initiate a sequence-specific double-strand break (DSB), after which the cell’s innate DNA repair mechanisms are triggered to engage in repair^7,8^. In the absence of a template to guide the repair, the DSB can be resolved by joining the two ends together through nonhomologous end-joining (NHEJ)^9–11^. This mechanism frequently incorporates base pair (bp) insertions or deletions (indels), that can disrupt the gene’s coding region and as such has been leveraged to effectively disrupt gene function^12^. Two alternative DSB repair pathways routinely exploited for precise gene insertions are: homology-directed repair (HDR) and microhomology-mediated end joining (MMEJ). Insertions using HDR requires a donor DNA template containing long stretches of homologous sequence flanking the intended modification site to mediate repair^13^. In contrast, MMEJ-based gene editing relies on the binding of microhomologies of ∼5-25 bp of complementary sequence to engage repair^14,15^. Employing these pathways improves editing accuracy over NHEJ, albeit at the cost of diminished efficiency, since these pathways exhibit lower activity levels and are only activated during specific stages of the cell cycle^16^. From an execution standpoint, they also necessitate the supply of loci-specific homologous sequences which can be difficult to amplify and integrate into repair constructs. Collectively, when attempting to insert large transgenes, these limitations often result in extremely low integration rates, making the isolation of homozygous cell lines both challenging and time-consuming^17–20^.

Instead of enhancing CRISPR or delivery components directly to address low integration rates, alternative approaches have long explored optimizing the donor construct itself to improve editing outcomes. This typically involves partitioning the donor cassette to include the desired tag or modification, along with a fluorescent or mammalian selectable marker to allow for easy enrichment of edited cells. Several methods have explored this concept^14,17,18,21–25^, with donor designs being largely summarized into two groups: promoter-based and multicistronic repair cassettes. Promoter-based cassettes involve the insertion of a tag within the reading frame of a GOI followed by a distinct promoter sequence that drives the expression of the selectable marker. In contrast, multicistronic approaches make use of sequence elements like internal ribosome entry sites (IRES)^26,27^ or 2A “self-cleaving” peptide sequences^28,29^ to facilitate efficient co-expression of the selectable marker. However, current tagging strategies are typically not easily amenable to switching tags or selectable markers and involve intricate cloning steps, making them less attractive for many labs to adopt.

Here, we’ve developed a set of optimized repair cassettes aimed at simplifying the process of editing and selecting for endogenously tagged cell lines. Our “quickTAG” (qTAG) system provides a plasmid scaffold that can be implemented with straight forward cloning approaches and can be easily adapted to accommodate alternative tags and selection markers. This study details the design and utility of these plasmids, showcasing their ability to facilitate both fluorescent and non-fluorescent tagging. Additionally, we provide insights into the timelines and strategies for efficiently generating homozygous cell lines and enabling multiple rounds of editing within the same line. Finally, we demonstrate the versatility of these cassettes through alternative editing scenarios, expanding their applications beyond traditional gene tagging to include conditional expression tagging, selectable knock-out tagging, and safe-harbour targeted expression. By providing convenient access to the qTAG library of plasmids through Addgene (https://www.addgene.org/Laurence_Pelletier/), our goal is to streamline the community’s adoption of endogenous tagging and encourage the adaptation of this plasmid library for future editing strategies and applications.

## Results

### A versatile scaffold to streamline endogenous tagging

The qTAG cassettes are built upon the core 2A-based cassette structure described previously^14,24^. The initial cassettes were designed and synthesized as gene fragments with the 2A sequence situated between a tag and resistance gene. Due to its transcriptional linkage and short sequence length, the 2A-based approach was adopted over an IRES element or a promoter to co-express a selectable marker that would enrich for edited cells. Four core features were integrated into the cassette structure to ensure ease of use while maintaining adaptability (Fig. 1). These included: (1) Cassette designs specific to both the N- and C-terminus encoding regions, to facilitate tagging at either end of a gene. (2) To simplify cloning, specific cloning sequences were placed around the cassette. These sequences permit a uniform cloning approach regardless of the tag, resistance, or specific sequences associated with the GOI. These sequences were designed to integrate both long and short homology arms, facilitating HDR or MMEJ for precise homology-based repair. (3) To enhance cassette adaptability, unique restriction sites were placed next to each critical sequence element within the cassette. This design allows for easy alteration through restriction cloning and permits future customization. (4) Finally, lox sites flank the selectable marker gene to enable the removal of the marker after transient introduction of the Cre recombinase^30,31^. The mutant lox sites lox66 and lox71 were chosen to ensure irreversible deletion of the selectable marker gene and recovery of selection sensitivity^32^. This is particularly useful in experimental conditions where only a single selectable marker can be used to sequentially tag genes.

**Figure 1.**
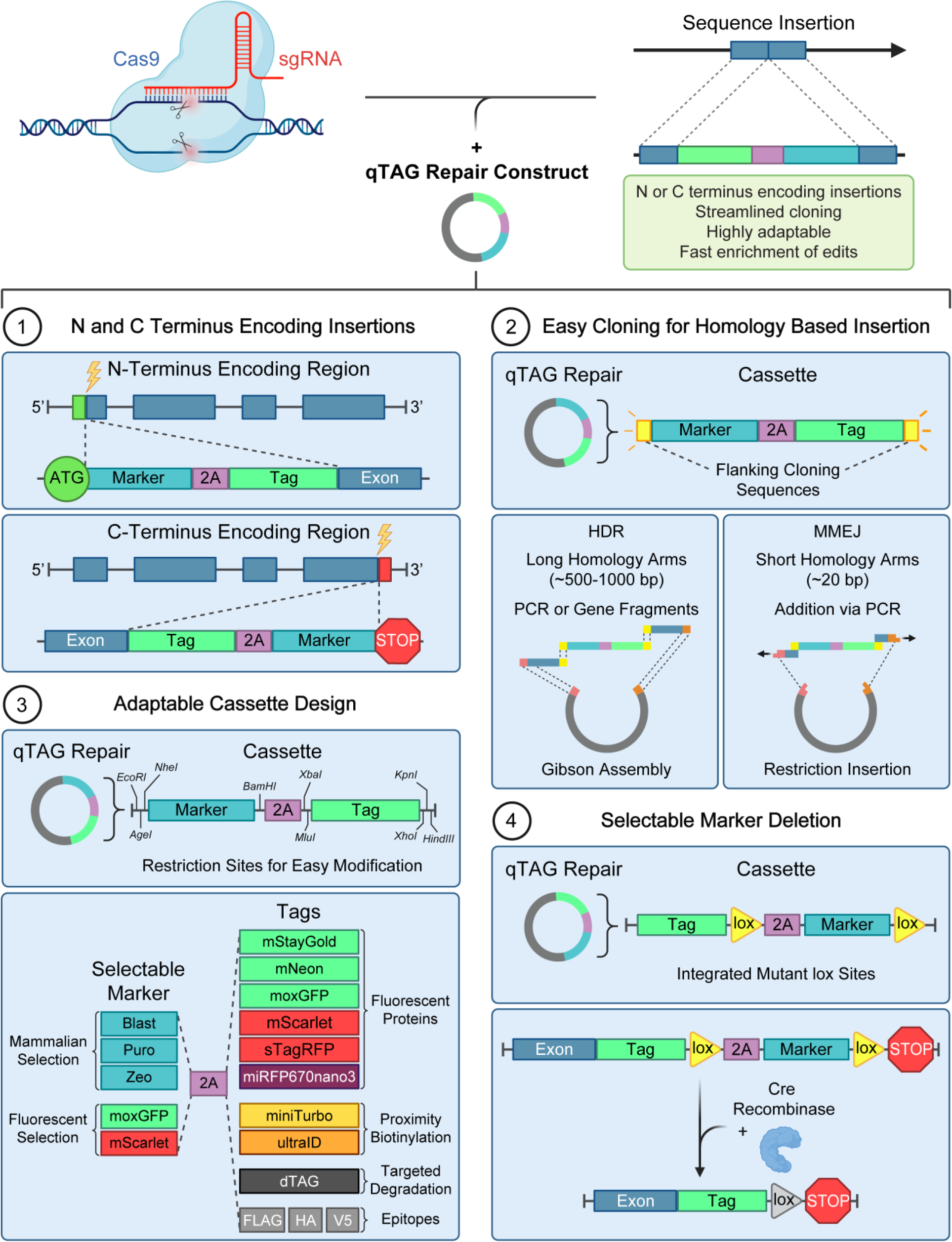
Features of qTAG cassettes. Four key features of the qTAG cassettes are depicted here. (1) 5’ and 3’ designs for N- and C-terminal modification; (2) the incorporation of uniform cloning sequences to streamline the cloning of long or short homology arms for HDR or MMEJ-based repair; (3) the inclusion of restriction sites near each genetic element in the cassette to conveniently swap tags or selectable markers; (4) the addition of lox sites on either side of the selectable marker to enable the marker gene removal.

The standardized cassette layout includes a selection of state-of-the-art tags to support a diverse array of assays. A panel of the brightest currently known monomeric fluorescent proteins covering the green, red, and far-red emission spectra enables observation and tracking of a target protein’s endogenous localization through live or fixed cell imaging. These include mStayGold^33^, mNeon^34^, moxGFP^35^, super-TagRFP^36^, mScarlet^37^, and miRFP670nano3^38^. The qTAG cassettes have also been expanded to support a wide variety of non-fluorescent tags. Proximity-dependent biotinylation enzymes, such as miniTurbo and ultraID, can be used to map interactions that occur in close physical proximity of a target protein^39,40^. For targeted-degradation studies, dTAG can be employed to induce rapid and reversible protein depletion with high specificity^24^. Finally, epitope tagging offers a solution in situations where antibodies against a protein of interest are unavailable, ineffective, or cross-reacts with paralogs. For this, we generated tagging cassettes containing 3xFLAG, 3xHA, and V5 epitopes, for which high-quality reagents are available. The use of these cassettes is further supported with a step-by-step protocol found here: https://www.biorxiv.org/content/10.1101/2023.11.01.565029v2.

### Editing and enrichment of fluorescent knock-ins with qTAG cassettes

To assess the functionality of the cassette design and the effectiveness of the editing and enrichment strategy, we initially chose to tag the histone protein H2BC11 with the fluorescent protein moxGFP (Fig. 2). The C-terminal region of H2BC11, was targeted using an sgRNA to permit the integration of a moxGFP-Puro cassette prior to the stop codon (Fig. 2a). We constructed moxGFP-Puro qTAG repair constructs containing both long and short homology arms to demonstrate the utility of both HDR and MMEJ for insertion and enrichment in HEK293T cells (Fig. 2b). When surveying cells for green fluorescence as an indication of cassette integration, a small proportion of cells in the initial pool expressing the H2B-moxGFP fusion was observed when tagged using either method (Fig. 2c,d). After selection with puromycin, a strong enrichment was observed in populations edited via HDR and MMEJ, as indicated by the shift in the distribution of cells displaying green fluorescence. We validated the target integration using fluorescence imaging to observe the localization of the endogenous fusion protein. Nuclear localization of H2BC11-moxGFP edited cells was confirmed through counterstaining with DAPI (Fig. 2e). Additionally, the presence of tagged alleles was verified by PCR using primers that anneal outside the homology arms. We observed a band shift consistent with the increased size of the qTAG-moxGFP-Puro cassette compared to the untagged allele (Fig. 2f). This tagging methodology was tested across various human cell lines including HAP1, ARPE-19, and U-2OS cells (Supp. Fig. 1). Despite differing initial tagging rates, consistent enrichment of edited alleles via HDR and MMEJ was observed post-selection with puromycin (Supp. Fig. 1a-d). The evaluation of alternative mammalian selection markers using blasticidin and zeocin also demonstrated their equivalence in enriching cells with tagged alleles (Supp. Fig. 2a,b). As a final evaluation using fluorescent tagging, we assessed the ability to introduce these insertions into H9 human embryonic stem cells. Repair constructs were devised to insert a qTAG-Blast-mScarlet cassette into the N-terminus encoding region of the tubulin gene TUBA1B. For editing, electroporation with precomplexed RNPs containing purified SpCas9 and an in-vitro transcribed sgRNA was used for delivery (Supp. Fig. 3a). Following delivery, selection-enrichment, and clonal expansion, mScarlet-TUBA1B expressing H9 cells were observed while still maintaining their pluripotent state, confirmed with staining against OCT4 (Supp. Fig. 3b).

**Figure 2.**
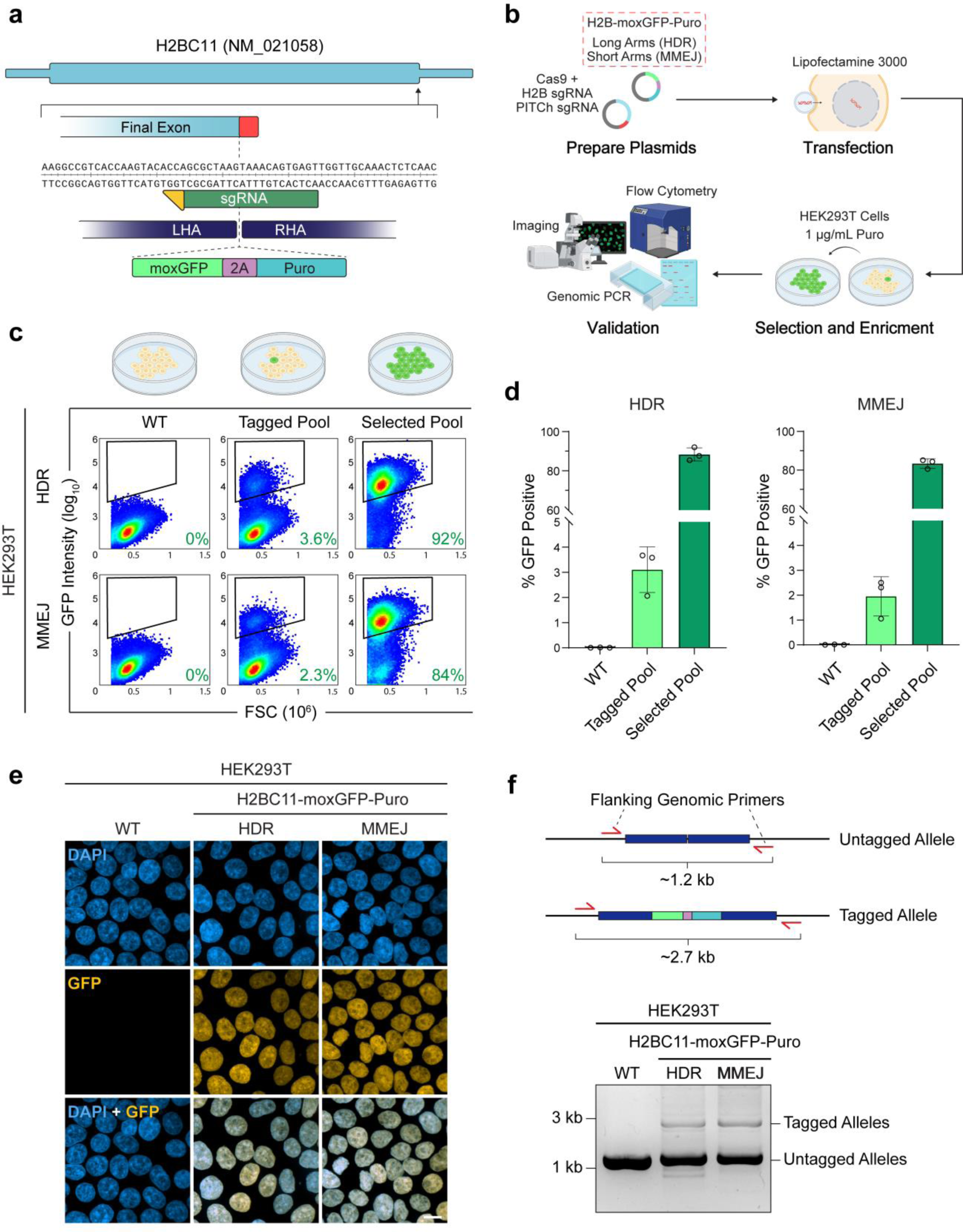
Editing, enrichment, and validation of fluorescent knock-ins with qTAG cassettes. (a) Schematic gene design of the H2BC11 gene displaying the binding of the sgRNA (dark green) and its PAM binding site (yellow), homology arms, and 3’ targeted insertion site. (b) An overview ofthe strategy used to carry out the endogenous tagging of H2BC11 with a C-terminus encoding insertion of the qTAG-moxGFP-Puro cassette via HDR (long homology arms) and MMEJ (short homology arms). (c) Representative flow-cytometry plots depicting 200,000 cells from the following samples: untagged wild-type, the initial tagged H2BC11-GFP-Puro pool, and the final H2BC11-GFP-Puro selected pools of HEK293T cells obtained through both HDR and MMEJ mechanisms. Gates indicate the threshold of fluorescent green cells within the total population as an indicator of positive editing. (d) Flow cytometry quantifications of GFP-positive cells, based on three biological replicates with each measurement encompassing 200,000 cells. (e) Representative images of WT, HDR-edited H2BC11-moxGFP-Puro, and MMEJ-edited H2BC11-moxGFP-Puro HEK293T cells co-stained with DAPI. (f) An agarose gel displaying the bands produced by genomic PCR with primers located outside the homology arms, probing for locus-specific integration of the qTAG cassette in HEK293T cells. Scale bars: 10 µm.

### Selectable marker gene excision and serial gene tagging

Next, we sought to demonstrate the ability to recover antibiotic sensitivity and subsequently use a single mammalian resistance gene for serial gene-tagging. To do this, we targeted two genes: the tubulin protein TUBB4B with a C-terminus encoding insertion of a mNeon-Blast cassette and the histone protein H3C2 with a C-terminus encoding insertion of a miRFP670nano3-Blast cassette (Fig. 3a). Editing commenced with TUBB4B in ARPE-19 cells. After blasticidin selection, the pool was transiently transfected with a plasmid encoding Cre-2A-Puro to excise the blasticidin resistance gene. Following a brief puromycin selection for two days to enrich transiently transfected cells, an enriched pool of TUBB4B-Neon cells expressing the Cre recombinase was generated. This pool was subsequently seeded for clonal expansion and a clonal homozygous cell line with a rescued sensitivity to blasticidin was isolated (Fig. 3b). PCR across the insertion junctions from cell lines representing the various phases of cell-line editing confirmed the presence of tagged alleles at the TUBB4B locus in the initial tagged pool (Fig 3c). A molecular weight band shift consistent with the loss of the blasticidin sequence was noticed in the Cre-transfected pool as well as the clonal cell line suggesting successful excision of the blasticidin gene. To further verify that the reading frame of TUBB4B was not altered, the TUBB4B locus was amplified from the TUBB4B-mNeon clonal cell line and Sanger sequenced. Due to the use of mutant lox sites, if the sequence between these elements was excised correctly, a mutant low-binding lox71/66 site would form. We amplified and sequenced the TUBB4B locus from the TUBB4B-mNeon clonal cell line and confirmed the formation of the correct low-binding lox sequence. The restored sensitivity to blasticidin after excision was then demonstrated in both the Cre-transfected pool and the resulting clonal lines by using selection assays and staining with crystal violet (Fig. 3d). Fluorescent imaging further confirmed the expected localization of the TUBB4B-mNeon endogenous protein across each of the cell lines generated during the editing phases (Fig. 3e). The renewed sensitivity of the APRE-19 TUBB4B-mNeon line was exploited to subsequently tag the histone protein H3C2 with miRFP670nano3-Blast (Fig. 3f,g).

**Figure 3.**
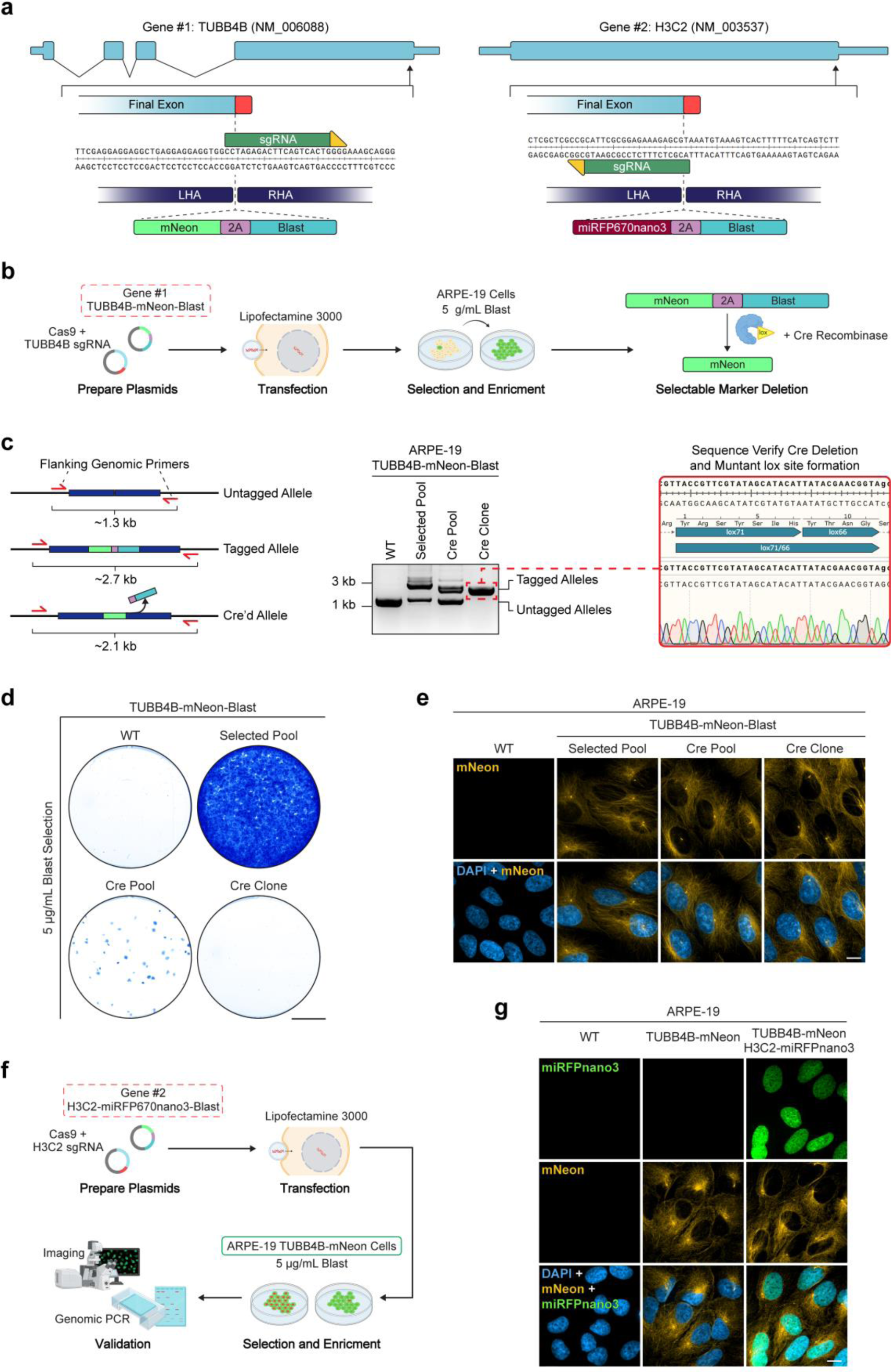
Selectable marker gene excision via the Cre recombinase and serial endogenous tagging. (a) Schematic gene designs of TUBB4B and H3C2. Shown are the location of the sgRNA (dark green) and its PAM binding site (yellow), homology arms, and targeted insertion site. (b) Overview of the strategy used to carry out the tagging of TUBB4B with a C-terminus encoding insertion of a qTAG-mNeon-Blast cassette. (c) An agarose gel displaying bands produced by genomic PCR with primers located outside the homology arms, probing for locus-specific integration of the qTAG cassette and Cre-edited alleles in ARPE-19 cells. Sanger sequencing of a homozygous clonal line displayed accurate excision of the marker gene as displayed by the formation of the inactive lox mutant site post Cre-editing. (d) Representative images of crystal violet-stained plate wells containing WT, selected pool, Cre pool, and Cre clonal cell lines of TUBB4B-mNeon APRE-19 cells selected with blasticidin. Scale bars: 1 cm. (e) Representative images of WT, selected pool, Cre pool, and Cre clonal cell lines of TUBB4B-mNeon APRE-19 cells co-stained with DAPI. Scale bars: 10 µm. (f) Overview of the strategy to carry out subsequent tagging of H3C2 with a C-terminus encoding insertion of qTAG-miRFP670nano3-Blast within the previously edited TUBB4B-mNeon ARPE-19 cells. (g) Representative images of WT, tagged TUBB4B-mNeon, and dual tagged TUBB4B-mNeon and H3C2-miRFP670nano3 ARPE-19 cells co-stained with DAPI. Scale bars: 10 µm.

### Non-fluorescent tagging applications

Examples of gene-tagging with qTAG cassettes containing functional proteomic tags are also presented here. The combination of low editing efficiency and the absence of fluorescent sorting options makes isolating insertions using these tags more challenging. However, coupling them with a selectable marker ensures that the process remains as efficient as our prior demonstrations of fluorescent protein insertion and enrichment. Here, the tagging scheme of the nuclear membrane protein LMNB1 is established and validated using various non-fluorescent tags, including the proximity biotin ligases miniTurbo and ultraID, the targeted degradation tag dTAG, and epitope tags FLAG, HA, and V5 (Fig. 4a). With HAP1 cells, the experimental scheme involving editing, selection, Cre-excision, and clonal expansion was followed to generate isogenic tagged lines for proteomic evaluation (Fig. 4b). First, the functionality of two biotin ligases (miniTurbo and ultraID) fused to endogenous LMNB1 was assessed for correct targeting and biotinylation activity. Imaging of HAP1 cells expressing V5-miniTurbo-LMNB1 and V5-ultraID-LMNB1 revealed robust nuclear membrane labelling after treatment with biotin and detection via streptavidin after 1 hour (Fig. 4c). Immunoblots using a streptavidin antibody and antibodies against V5 and LMNB1 further confirmed robust biotinylation of proximal proteins within just 10 minutes, with even greater biotinylation detected after 1 and 12 hours (Fig. 4d). Imaging of HAP1 cells expressing V5-dTAG-LMNB1 and co-stained with anti-V5 and anti-LMNB1 antibodies confirmed nuclear membrane localization (Fig. 4e) that was reduced in cells after treatment with dTAG-13, the small molecule responsible for inducing degradation17. Immunoblotting also confirmed the time-dependent degradation of V5-dTAG-LMNB1 in cells treated with dTAG-13 (Fig. 4f). Lastly, for epitope tagging, the general epitopes FLAG, HA, and V5 were used to tag LMNB1. Immunofluorescence of HAP1 cells stained with epitope-specific antibodies revealed nuclear membrane localization of 3xFLAG-LMNB1, 3xHA-LMNB1, and V5-LMNB1 (Fig. 4g). Additionally, immunoblotting with epitope-specific and target-specific antibodies resulted in robust detection of the endogenous fusion protein, confirming the proper integration of the cassettes and localization of the endogenous fusion (Fig. 4h).

**Figure 4.**
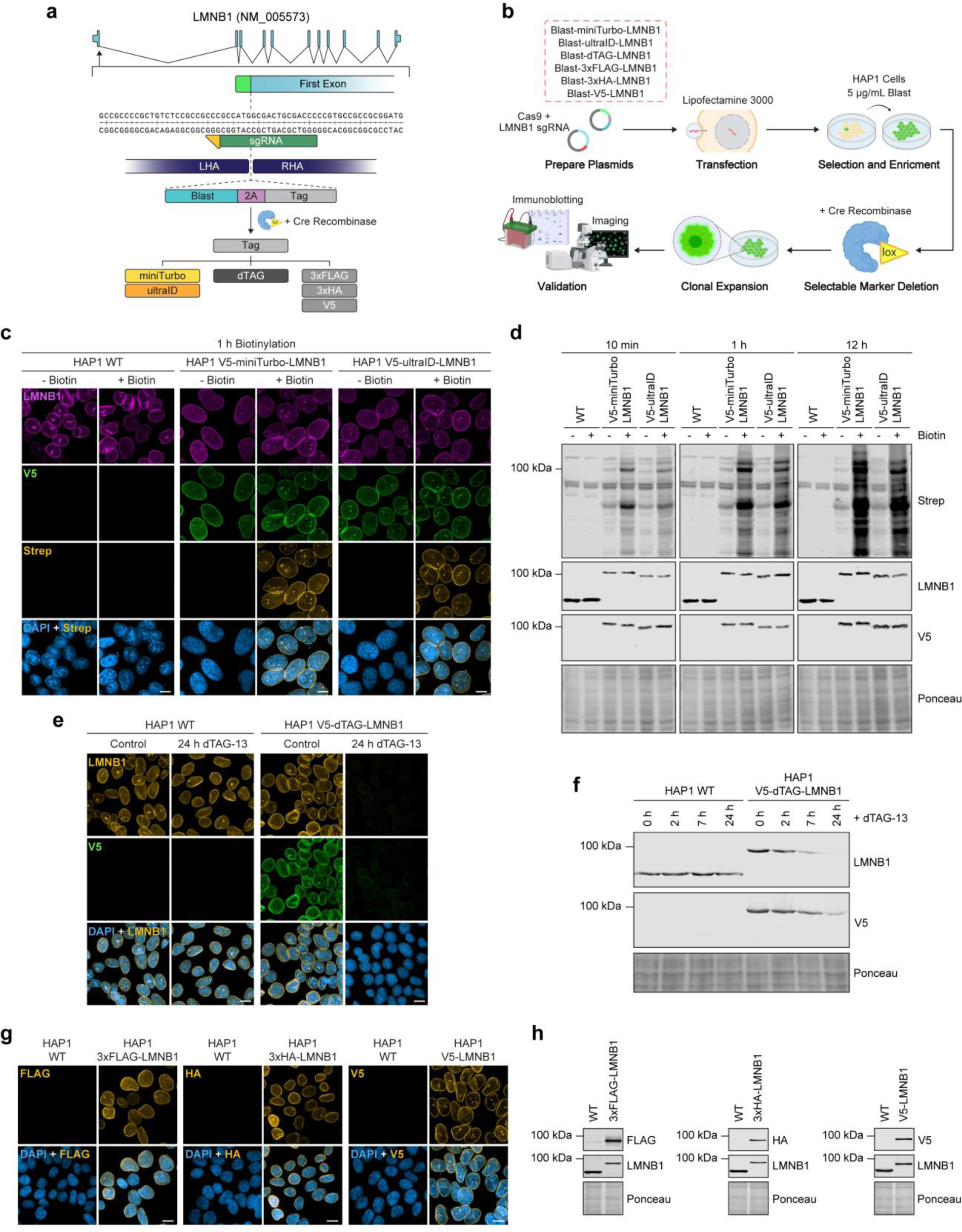
Non-fluorescent gene qTAG applications and validation. (a) Schematic gene design of LMNB1 displaying the binding of the sgRNA (dark green) and its PAM binding site (yellow), homology arms, and targeted insertion site. (b) Overview of the strategy to carry out tagging of LMNB1 with a variety of functional proteomic qTAG cassettes. (c) Representative images of WT, V5-miniTurbo, and V5-ultraID tagged LMNB1 in HAP1 cells. Cells were stained with DAPI and probed with streptavidin in the absence or presence of biotin for 1 hour. Scale bars: 10 µm. (d) A representative immunoblot of WT, V5-miniTurbo, and V5-ultraID tagged LMNB1 in HAP1 cells probed with antibodies against streptavidin, V5, and LMNB1 in the absence or presence of biotynilation for 10 min, 1h, and 12h. (e) Representative images of WT and V5-dTAG tagged LMNB1 in HAP1 cells stained for DAPI and probed for V5 in the absence or presence of dTAG-13 for 24h. Scale bars: 10 µm. (f) A representative immunoblot of dTAG-V5 tagged LMNB1 in HAP1 cells probed for V5 and LMNB1 in the absence or presence of dTAG-13 for 2, 7 and 24h. (g) Representative images of WT, 3xFLAG tagged, 3xHA tagged, and V5 tagged LMNB1 in HAP1 cells stained for DAPI and probed for their respective epitopes. Scale bars: 10 µm. (h) Representative immunoblots of WT, 3xFLAG tagged, 3xHA tagged, and V5 tagged LMNB1 in HAP1 cells with their respective epitope-specific antibodies.

### Compatibility of the qTAG scaffold for the integration of novel tags

We next sought to showcase the adaptability of the qTAG cassette system for future applications by incorporating novel tags. StayGold is a bright green fluorescent protein published in 2022 and is known for its exceptional photostability, which far surpasses that of its contemporary fluorescent proteins^41^. However, its inclination to form an obligate dimer to function poses a hindrance in protein fusion applications. Recently, monomeric variants of StayGold were designed to overcome these challenges including mStayGold^33^, StayGold-E138D^42^, and mBaoJin^43^. This presents an opportunity for gene tagging applications that facilitate visualization of endogenously tagged proteins, including those expressed at low levels, in long-term live cell imaging. Due to its enhanced brightness, we opted to pursue a Tag-Only enrichment scheme. In plug-and-play fashion, leveraging the restriction sites within the qTAG cassette, we initiated the process by subcloning the mStayGold sequence into the N- and C-terminus targeting qTAG cassettes (Fig. 5a). Homology arms were subsequently incorporated, and genes tagged with mStayGold were enriched through fluorescence cell sorting. To showcase mStayGold’s photostability, the actin gene ACTB was tagged with both mNeon and mStayGold in RPE-1 PAC (-) cells. Homozygous clones of each cell line underwent live-cell imaging with continuous exposure where the normalized fluorescence intensity could be quantified over time (Fig. 5B). Imaging and intensity measurements of these native fusions revealed mStayGold’s resilience against photobleaching, exhibiting a discernible difference in intensity within two minutes of acquisition, when compared to mNeon-ACTB, and further maintaining its photostability throughout the entire acquisition period. Tas an extension of this application, mStayGold-ACTB cells were subjected to live-cell timelapse super-resolution imaging (Fig. 5C). Super-resolution imaging is typically more challenging due to its higher light load, often limiting its application to fixed cell imaging. When combined with endogenous tagging using traditional fluorescent proteins like GFP or mNeon, this challenge is exacerbated, resulting in insufficient signal over time to generate a nice movie. Yet, the enhanced brightness and photostability of mStayGold can reconcile these challenges, allowing for super-resolution structured illumination time-lapse microscopy where the leading-edge dynamics in mStayGold-ACTB RPE-1 PAC (-) cells can be observed. Finally, to demonstrate the versatility of mStayGold in endogenous molecular fusions, mStayGold qTAG cassettes were used to target genes that illuminate a diverse range of cellular structures (Fig. 5D). There included high-abundance, polymerizing proteins like α-Tubulin to low-abundance proteins localized at the centrosome.

**Figure 5.**
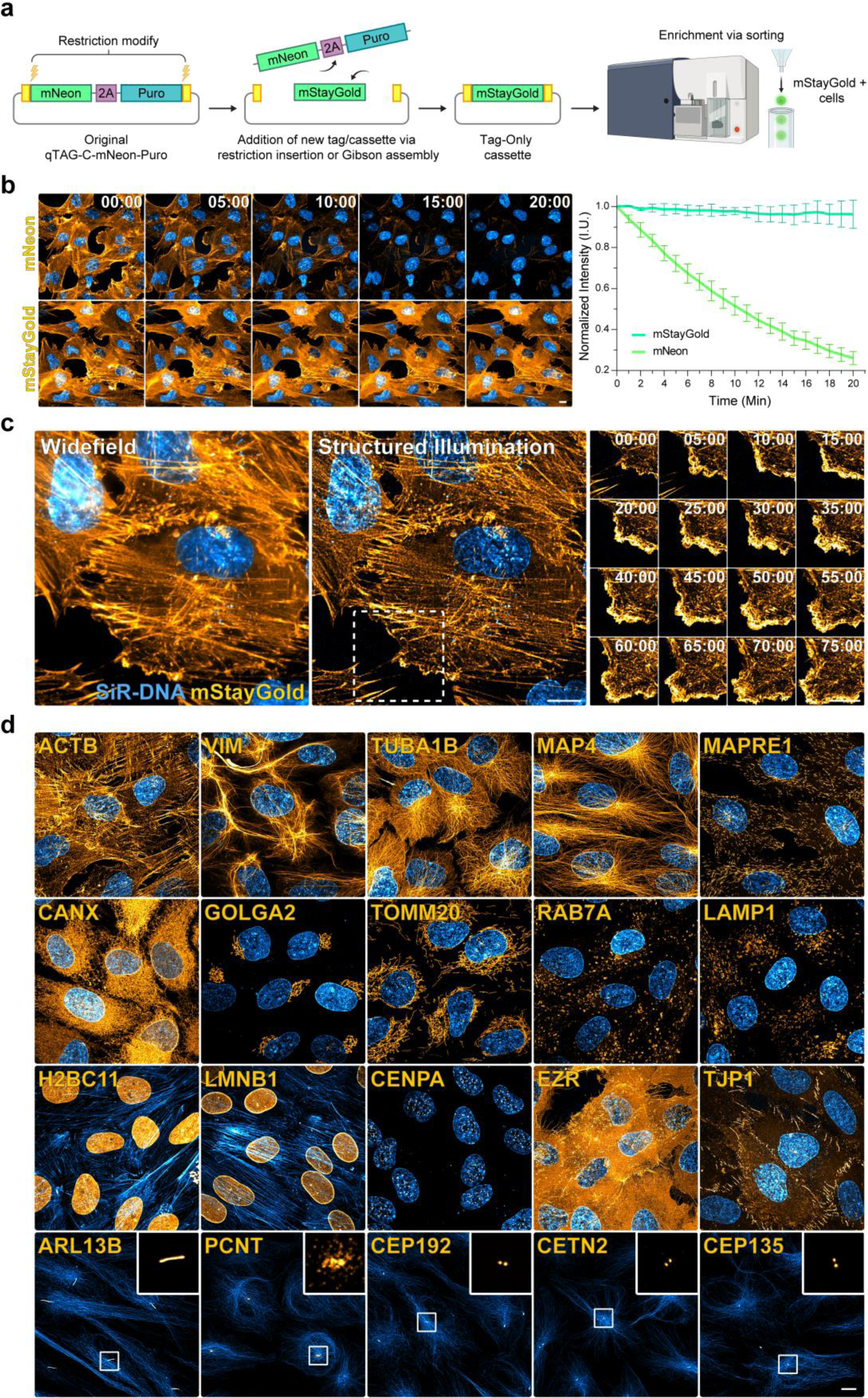
Integration and capability of mStayGold in qTAG tagging cassettes. (a) A schematic of the modification process to include mStayGold in a Tag-Only knock-in cassette followed by a cell sorting enrichment strategy. (b) Left - Representative live images of N-terminally tagged mNeon-ACTB and mStayGold-ACTB in RPE-1 cells. Cells were live co-stained with SiR-DNA and imaged continuously for 20 min. (b) Right – Normalized intensity measurements of each cell line from three biological replicates, each comprised of four technical replicates. (c) Super-resolution timelapse imaging of mStayGold tagged ACTB co-stained with SiR-DNA. (c) Left – A widefield image at the start of the timelapse. (c) Middle – The corresponding structured illumination super-resolution reconstructed image. (c) Right – A time-lapse array of images following the highlighted area.(d). Endogenous molecular fusions with mStayGold. Representative images mStayGold (Orange) tagged to genes highlighting various subcellular compartments, co-stained with SiR (Blue) dyes against Actin, Tubulin, or DNA. Scale bars: 10 µm.

### Exploring different editing applications by modifying the core qTAG cassette structure

Beyond the core qTAG cassette framework, researchers possess the freedom to modify the insertion cargo sequence in creative ways that can allow for alternate enrichment schemes or new tagging scenarios altogether. Here we present three scenarios as examples that expand the utility and applications of the qTAG scaffold.

In situations where the target gene exhibits low or no expression, for example when gene expression fluctuates during differentiation or reprogramming, accumulating a sufficient amount of selectable marker to confer resistance for the enrichment for integrations can be challenging. The tagging of these genes can be achieved by replacing the “2A” element with the constitutive promoter such as the short form of the EF1a promoter (EFs) (Fig. 6A).

**Figure 6.**
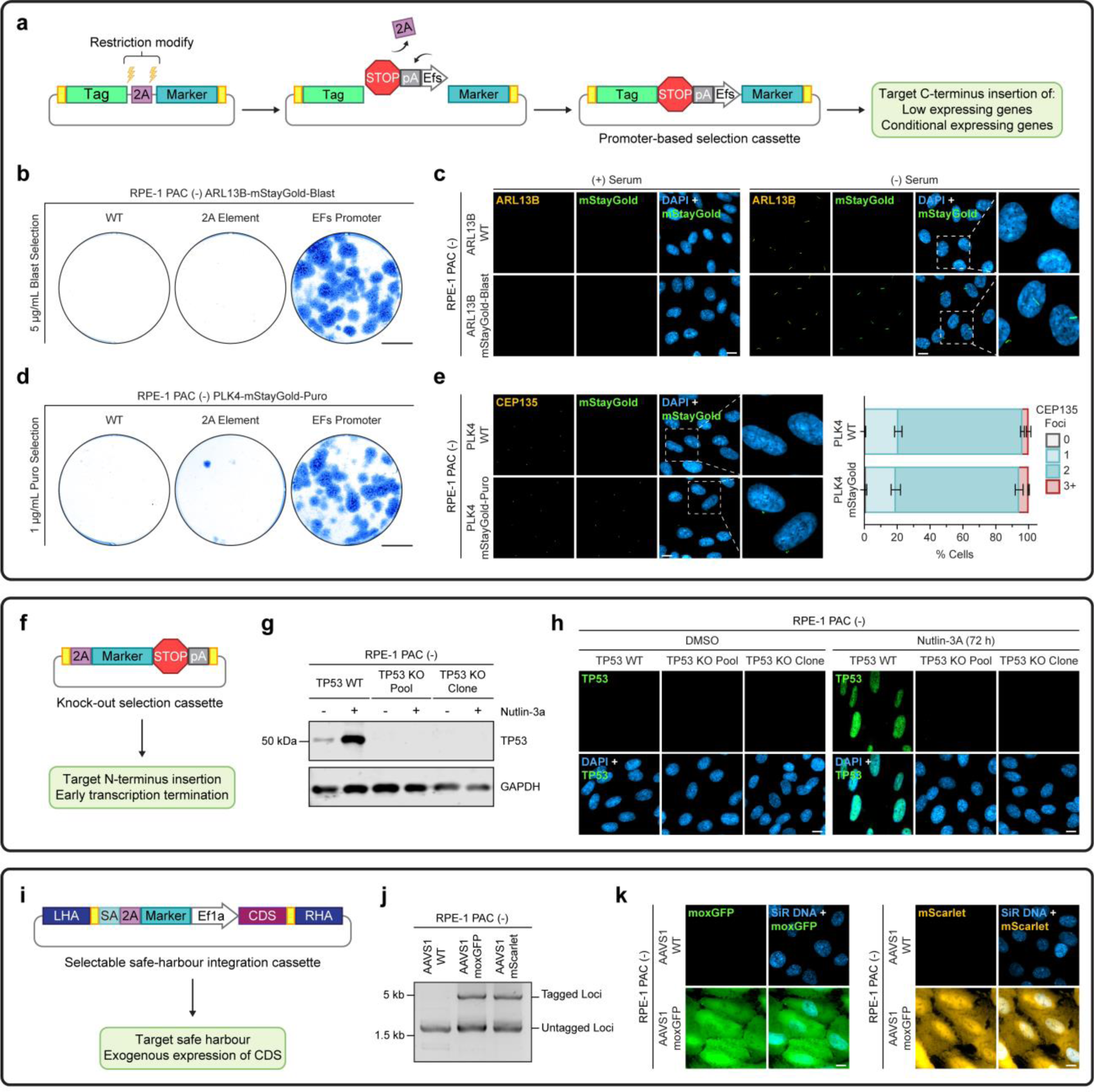
Generating alternative cassette designs for use cases beyond endogenous tagging. (a) A schematic of the modification process to convert the cassette from a multicistronic 2A-based approach to a promoter-based approach for the enrichment of integrations. pA = polyadenylation signal, EFs = EF1a short promoter. (b) Representative images of crystal violet-stained plate wells containing WT-ARL13B, initially tagged 2A-based ARL13B-mStayGold, and initially tagged promoter-based ARL13B-mStayGold RPE-1 PAC (-) cells selected with blasticidin for 7 days. Scale bars: 1 cm (c) Representative images of WT and endogenously tagged, promoter-based ARL13B-mStayGold. Cells were stained with DAPI and probed with anti-ARL13B in the absence or presence of serum starvation for 72 h. Scale bars: 10 µm. (d) Representative images of crystal violet-stained plate wells containing WT-PLK4, initially tagged 2A-based PLK4-mStayGold, and initially tagged promoter-based PLK4-mStayGold RPE-1 PAC (-) cells selected with puromycin for 7 days. Scale bars: 1 cm. (e) Left – Representative images of WT and endogenously tagged, promoter-based PLK4-mStayGold. Cells were stained with DAPI and probed with anti-CEP135. Scale bars: 10 µm. (e) Right – a quantification of centrosome number within the WT and promoter-based PLK4-mStayGold pools. >100 cells were quantified in each of the 3 biological replicates in each condition. (f) – A schematic of a cassette that expresses a selectable marker followed by the induction of transcriptional termination allowing for selectable knockouts if integrated at the N-terminal encoding region of a gene. (g) A representative immunoblot of RPE-1 PAC (-) WT-TP53 cells, a TP53-KO-Puro selected pool of cells, and a TP53-KO-Puro selected clonal cell-line probed with antibodies against TP53 and GAPDH in the absence or presence of 10µM Nutlin-3a for 72 h. (h). Representative images of RPE-1 PAC (-) WT-TP53 cells, a TP53-KO-Puro selected pool of cells, and a TP53-KO-Puro selected clonal cell-line probed with anti-TP53 in the absence or presence of 10µM Nutlin-3a for 72 h. (i). A schematic of a cassette that allows for the integration at the safe-harbour AAVS1 loci, allowing for exogenous expression. SA = splice acceptor, CDS = coding sequence. (j). Genomic PCR outside the homology arms probing for locus-specific integration of the AAVS1 safe-harbour cassettes expressing exogenous moxGFP and mScarlet in RPE-1 PAC (-) cells. (k) Representative images of RPE-1 PAC (-) WT-AAVS1 cells, a AAVS1-moxGFP puro selected pool of cells, and a AAVS1-mScarlet puro selected pool of cells co-stained with DAPI. Scale bars: 10 µm.

In our testing, when targeting the cilia gene ARL13B and the centrosome duplication gene PLK4 with a mStayGold-2A-based cassettes, selection assays revealed low to no resistant colonies post-selection and crystal violet staining (Fig. 6B,D). However, when targeting these same genes with mStayGold cassettes with EFS produced selectable marker, many resistant colonies were observed after selection. Imaging of ARL13B-mStayGold-Blast cells in serum supplemented media and after 72 h of serum starvation confirmed the formation and localization of ARL13B-mStayGold to the cilia when co-stained with an antibody against ARL13B (Fig. 6C - Left). The imaging of PLK4-mStayGold-Puro cells also confirms the localization of PLK4-mStayGold to the centrosomes when co-stained with ant-CEP135, a marker for centrioles (Fig. 6E). PLK4 regulates centriole duplication, and its overexpression is known to induce the formation of extra centrosomes^44,45^. Thus, by co-staining with anti-CEP135, we could enumerate the centrosomes to determine if extra centrosomes occurred in our edited line. Quantification of centrosome numbers in both our RPE-1 PAC (-) background and our edited PLK4-mStayGold cell line revealed no significant difference between populations with normal centrosome numbers (cells with 1-2 CEP135 foci) and those with amplified centrosome numbers (cells with 3 or more CEP135 foci). This confirms that the endogenous fusion did not alter PLK4’s duplication function, thereby enabling its native study (Fig. 6C - Right).

Current methods of creating gene disruptions rely on the infidelity of NHEJ repair pathway to create random insertions or deletions that disrupt the reading frame. As such, clonal selection and verification is required. Alternatively, a drug selection cassette inserted at the N-terminus encoding region of a gene to rapidly select for targeted gene knockouts. Here, the qTAG cassette was customized to include a selectable marker terminated with a stop codon, followed by transcriptional termination sequence in the form of the SV40 polyadenylation signal (see Fig. 6F). To showcase the effectiveness of this cassette, we targeted the tumor suppressor protein TP53 where editing, selection enrichment, and homozygous clones were isolated in RPE-1 PAC (-) cells. Cells were then treated with Nutlin-3a, a small molecule inhibitor of MDM2, to trigger a TP53-mediated response that would lead to the accumulation of TP53 in TP53-WT cells^46^. Immunoblotting of TP53-WT, a selected pool of edited TP53-KO-Puro cells, and a clonally edited TP53-KO-Puro line with an antibody against TP53, in the presence or absence of Nutlin-3A, had displayed the accumulation of TP53 only in the treated TP53-WT scenario (Fig. 6G). The selected pool and clonal cell line had both exhibited ablation of TP53 signal in both treatment conditions. This phenomenon was further confirmed via immunofluorescence (Fig. 6H).

Finally, safe harbour sites refer to loci in the genome that are capable of accommodating the integration of new genetic elements, ensuring their predictable functionality while minimizing the risk of disrupting normal cellular processes. The AAVS1 safe harbour site was selected to incorporate a modified qTAG cassette for gene expression. This cassette removes the core tag-2A-marker structure and replaces it with a selectable marker, succeeded by a full length EF1a promoter and a multiple cloning site. This configuration enables the seamless integration of a coding sequence (CDS) of your preference (Fig. 6I). To showcase the functionality of this safe harbour expression cassette, two examples of moxGFP and mScarlet were cloned into the CDS location. The homology arms targeting AAVS1 were then integrated to enable widespread moxGFP and mScarlet expression within the cell upon integration. Following editing, genomic junction PCR on selected pools confirmed the insertion of the ∼4.3 kb expression cassette into the targeted safe-harbour region (Fig. 6J). Live-cell imaging of AAVS1-moxGFP and AAVS1-mScarlet selected cells stained with SiR-DNA confirmed the expression of moxGFP and mScarlet throughout the cells, respectively (Fig. 6K).

## Discussion

The study of protein function by ectopic expression is a mainstay of scientific experimentation. In an ideal world, protein behaviour would be observed without any modification, however this is not yet practical. Observing tagged proteins expressed at their endogenous levels is currently the best method to study protein function. The emergence of the CRISPR/Cas9 system has enabled the widespread use of endogenous tagging over traditional overexpression methods to explore gene function and protein behavior. Our main objective in developing this plasmid system was to expand the accessibility of this valuable approach. Harnessing the cell’s intrinsic repair mechanisms for precise gene editing is frequently inefficient, necessitating a significant amount of time to isolate homozygous cell lines. However, by optimizing the donor cassettes to include a linked selectable marker, the enrichment of edited cells can be easily achieved.

In scenarios focused on integrating functional tag sequences onto genes, we presented qTAG cassettes containing both fluorescent and non-fluorescent tags. Alongside the cassettes, we detail a strategy demonstrating how these constructs enable accurate editing and selection-based enrichment of edited cells. Additionally, the integration of a wide array of high-performance tags outlines their utility for both fluorescent and non-fluorescent functional assays in studying gene function. Several different strategies have been developed to address the challenges associated with endogenous tagging in human cells^14,17–19,21,23,47–50^. Each of these techniques offers different advantages and features with some being highly tailored for specific applications such as post-mitotic tagging^50^ or stem cell tagging^19^. These systems encompass several variations of the formula, including different Cas enzymes, formats of donor repairs, delivery methods, and post-editing enrichment schemes. While these variations offer benefits for specific scenarios, they also have limitations and differ in requirements for equipment, reagents, cloning capabilities, and editing timelines. Moreover, certain donor strategies have fixed designs that make it challenging to accommodate changes, such as using a different tag or selectable marker. In this context, our implementation sought to standardize and openly distribute a widely accessible framework, incorporating numerous principles from prior approaches. This initiative aimed to enable others to use, modify, and tailor the scaffold for their individual experiments.

The adaptability of these constructs is a major highlight of this system. We showcased the adaptability of this framework by incorporating new tags like mStayGold, which demonstrated its capability for long-term live cell fluorescence imaging of proteins at native levels. The simplicity of swapping selectable markers and tags with alternative or novel options will empower researchers to develop cassette variants beyond our current offerings. Expansion of proteomic tagging could encompass alternative degron tags, HaloTag, SnapTag, and ALFA tag, while fluorescent tags with emission spectra not covered in our collection (cyan, yellow, near-infrared) could be easily accommodated. Furthermore, this framework opens avenues for exploring the insertion of any cargo sequence within the “Tag” region, potentially facilitating the attachment of sequence elements such as signal peptides, subcellular targeting domains, cleavage signals, and binding domains. This allows for creative approaches to manipulate endogenous gene function or localization, that could provide deeper insights into their functionality.

However, within the standardized structure of the provided qTAG cassettes, two limitations do exist. One drawback is the reliance on transcriptional linkage of the target gene to the selectable marker with the 2A sequence. This requires that the target GOI be expressed at high enough levels in the cell to produce enough antibiotic resistance for efficient selection. Also, while 2A peptide sequences have garnered mass adoption in molecular biology to achieve co-translation, their cleavage or ‘skipping’ efficiency is not always consistent in all cellular backgrounds. This can result in a minor amount of read-through^28,51^ resulting in fusions between the tagged GOI and the selectable marker. Given the adaptable nature of the qTAG cassette system, we addressed these limitations providing separate cassettes where the tag region is terminated with a stop codon and the 2A sequence is substituted with an EFs promoter. This modification allowed the selection of edits independent of the target gene’s transcript levels and circumvents the constraints associated with 2A peptide sequences. Thanks to the constitutive expression of the selectable marker, even genes that are expressed at low levels or transcriptionally silent can be edited. This could be particularly useful in cases where the tracking of fate-based gene expression is important. As an example, during the differentiation or reprogramming of cells, the activation of fate-specific transcription factors often occurs. By silently tagging these genes at their C-terminus encoding region, either directly with a fluorescent protein sequence or with a 2A-fluorescent protein sequence, a cell line with an intrinsic disposition marker can be generated. Upon reaching the particular fate where the targeted gene would be expressed, the tagged transcription factor would fluoresce if directly tagged or illuminate the entire differentiating/reprogramming cell if tagged with a 2A-Fluorescent protein sequence, effectively serving as a transcriptional biosensor for cellular fate.

Beyond the core tag-2A-marker structure, we further modified the payload insertion region to explore two alternative tagging scenarios: facilitating selectable gene knockouts and enabling safe-haven exogenous expression. In the context of selectable knockout cassettes, there are situations where transfectability, poor editing efficiency, and/or clonal expansion is impractical due to limited replicative potential (such as in primary cells). In such instances, employing a selectable knockout scenario significantly shortens the timeline compared to traditional methods involving clonal indel validation. Although considerations regarding ploidy and copy number integration persist in both scenarios, the availability of a selectable enrichment option alleviates the challenges associated with factors like transfection efficiency and sgRNA binding efficiency. This was illustrated by the significant reduction in the target protein observed even within our selected pool of TP53-KO cells.

Our last alternative tagging approach focused on tagging the safe-harbour site AAVS1 with an expression cassette. This represents the most extensive alteration of the cassette structure, introducing multiple new genetic elements to fulfill a completely different purpose in accommodating exogenous expression. Common methods for achieving permanent integration of expression cassettes involve either random integration of a plasmid post-transfection or the use of lentivirus systems. However, these methods either depend on chance or entail specific considerations regarding viral culture, and result in random integration into the genome. Conversely, the preference often lies in integrating expression cassettes into predetermined genomic loci, a feat typically accomplished via the FLP-In system^52^. However, this process necessitates specialized FRT reagents, cell lines, and is time intensive. The safe harbor overexpression tagging approach addresses many of these challenges by facilitating quick, permanent, and site-specific integration for the expression of a target CDS.

In summary, we have detailed the development and application of a versatile endogenous tagging platform. Engineered with the aim of making CRISPR-mediated tagging more accessible, the qTAG library of plasmids incorporate various essential characteristics that enhance usability and adaptability, serving as an open framework for researchers to tailor towards their own experiments.

## Methods

### qTAG system construct development

The initial structures of the C- and N-terminus targeting cassettes were sourced from Nabet et al.^24^. These sequences were arranged from 5’ to 3’ to include a protein tag sequence in frame with a P2A peptide sequence, succeeded by a selectable marker sequence (with the tag and selectable marker sequence switched for the N-terminal targeting cassettes). Unique restriction sites were introduced next to every sequence element to allow for easy variant cloning. Prior to synthesizing codon-optimized gene fragments for the initial C- and N-terminus cassettes (GeneArt Strings, ThermoFisher), uniform cloning sequences were introduced around the cassette and lox sites to flank the selectable marker region. Following this, the cassettes were cloned into a pUC19 backbone. Variants of the cassette were produced via restriction insertion cloning, using specific enzymes tailored for the qTAG cassette and the insertion sequences (AgeI-HF, BamHI-HF, EcoRI-HF, HindIII-HF, KpnI-HF, MluI-HF, NheI-HF, XbaI, XmaI, NEB), followed by ligation (T4 DNA Ligase Kit, M0202, NEB) of the resulting products at a 1:3 molar ratio of vector to insert. Table 1 presents the sources of the tag sequences used. Separately, transient Cre + selectable marker expression plasmids were generated to complement the qTAG cassettes. Fragments containing an Ef1a promoter, Cre recombinase CDS, P2A element, puromycin selectable marker CDS, and a polyA signal were synthesized, followed by Gibson assembly (Gibson Assembly Master Mix, E2611L, NEB) at a 1:3 molar ratio of vector to fragment for directional cloning into a pUC19 backbone. All presently generated variants of qTAG cassettes and Cre plasmids are listed in Table 4.

**Table 1.** Sources of Tag Sequences.

| Tag | Source Plasmid | Source Study | Modifications |
| --- | --- | --- | --- |
| Cterm mStayGold | https://www.addgene.org/212019/ | (Ando et al.)<sup>33</sup> | Removal of internal restriction sites, codon optimization |
| Nterm mStayGold | https://www.addgene.org/212018/ | (Ando et al.)<sup>33</sup> | Removal of internal restriction sites, codon optimization |
| mNeon | https://www.addgene.org/58179/ | (Shaner et al.)<sup>34</sup> | Removal of internal restriction sites, codon optimization |
| moxGFP | https://www.addgene.org/68070/ | (Costantini et al.)<sup>35</sup> | Removal of internal restriction sites, codon optimization |
| mScarlet | https://www.addgene.org/85042/ | (Bindels et al.)<sup>37</sup> | Removal of internal restriction sites, codon optimization |
| super-TagRFP | https://www.addgene.org/173014/ | (Mo et al.)<sup>36</sup> | Removal of internal restriction sites, codon optimization |
| miRFP670nano3 | https://www.addgene.org/184664/ | (Oliynyk et al.)<sup>38</sup> | Removal of internal restriction sites, codon optimization |
| miniTurbo | https://www.addgene.org/107174/ | (Branon et al.)<sup>40</sup> | Removal of internal restriction sites, codon optimization |
| ultraID | https://www.addgene.org/172878/ | (Kubitz et al.)<sup>39</sup> | Removal of internal restriction sites, codon optimization |
| dTAG | https://www.addgene.org/91797/ | (Nabet et al.)<sup>24</sup> | Removal of internal restriction sites, codon optimization |
| 3xFLAG | Synthesized | - | - |
| 3xHA | Synthesized | - | - |
| V5 | Synthesized | - | - |

### Gene knock-in design, CRIPSR and repair construct assembly

CHOPCHOP v3^53^ (https://chopchop.cbu.uib.no/) was our primary sgRNA design tool because of its user-friendly graphical interface, which provides clear visualization of genomic loci and sgRNA binding sites. Selection of suitable sgRNAs for gene tagging design relied on several criteria. First, sgRNAs were locally filtered, being restricted to sgRNAs binding positions within ∼30bp of the 5’ or 3’ ends of a gene, depending on whether tagging was planned for the N- or C-terminus, respectively. Beyond location, predicted on-target and off-target scores were considered when choosing and ranking potential sgRNAs. To produce the sgRNAs within cells, we employed either plasmids encoding Cas9 and target sgRNA or delivered the Cas9 protein and sgRNAs directly as a ribonucleic protein complex (RNP). For plasmid-based delivery, the original Zhang protocol for SpCas9 plasmids (https://www.addgene.org/crispr/zhang/) was used to clone and deliver our Cas9 and sgRNA^54^. We particularly used either pX330 (https://www.addgene.org/42230/) or the MMEJ PITCh-primed variation of px330, the px330-PITCh plasmid (https://www.addgene.org/127875/)^14,25^. For RNP based delivery, Cas9 was expressed from a pET-based T7 promoter-containing plasmid (https://www.addgene.org/62934/) in BL21 bacteria^55^. The purification of Cas9 was carried out from a previously described method^56^. To produce the sgRNA, a one-step PCR assembly method^57^ was performed to generate the guide DNA basis. In-vitro transcription (HiScribe T7 High Yield RNA Synthesis Kit E2040L, NEB) from the DNA was then carried out, followed by RNA purification (RNAClean XP, Beckman Coulter).

For HDR-based qTAG repair constructs, 500-1000 bp of homologous genomic sequence flanking the predicted sgRNA cut site was either directly amplified from the genome via PCR or synthesized as gene fragments. The homology arms were generated with specific flanking cloning sequences outlined in Table 2 that are homologous to those on the qTAG cassettes. To ensure that the sgRNA sequence did not overlap with the donor construct, sgRNAs were chosen such that they would inherently disrupt the sgRNA sequence upon insertion. In situations where this was not feasible, PAM blocking or sgRNA blocking mutations were introduced during homology arm synthesis to prevent Cas9 cleavage of the donor construct upon entry into the nucleus or re-cleavage of the genomic insertion post-integration. For the final two fragments, the backbone fragment involved digesting the pUC19 backbone using EcoRI and HindIII enzymes, whereas the designated qTAG cassette intended for insertion was amplified using the cassette-specific primers listed in Table 2. These four fragments of digested pUC19 backbone, left homology arm, amplified qTAG cassette, and right homology arm were then directionally cloned using Gibson assembly at a 1:3:3:3 molar ratio to form the final qTAG HDR-based repair construct.

**Table 2.**
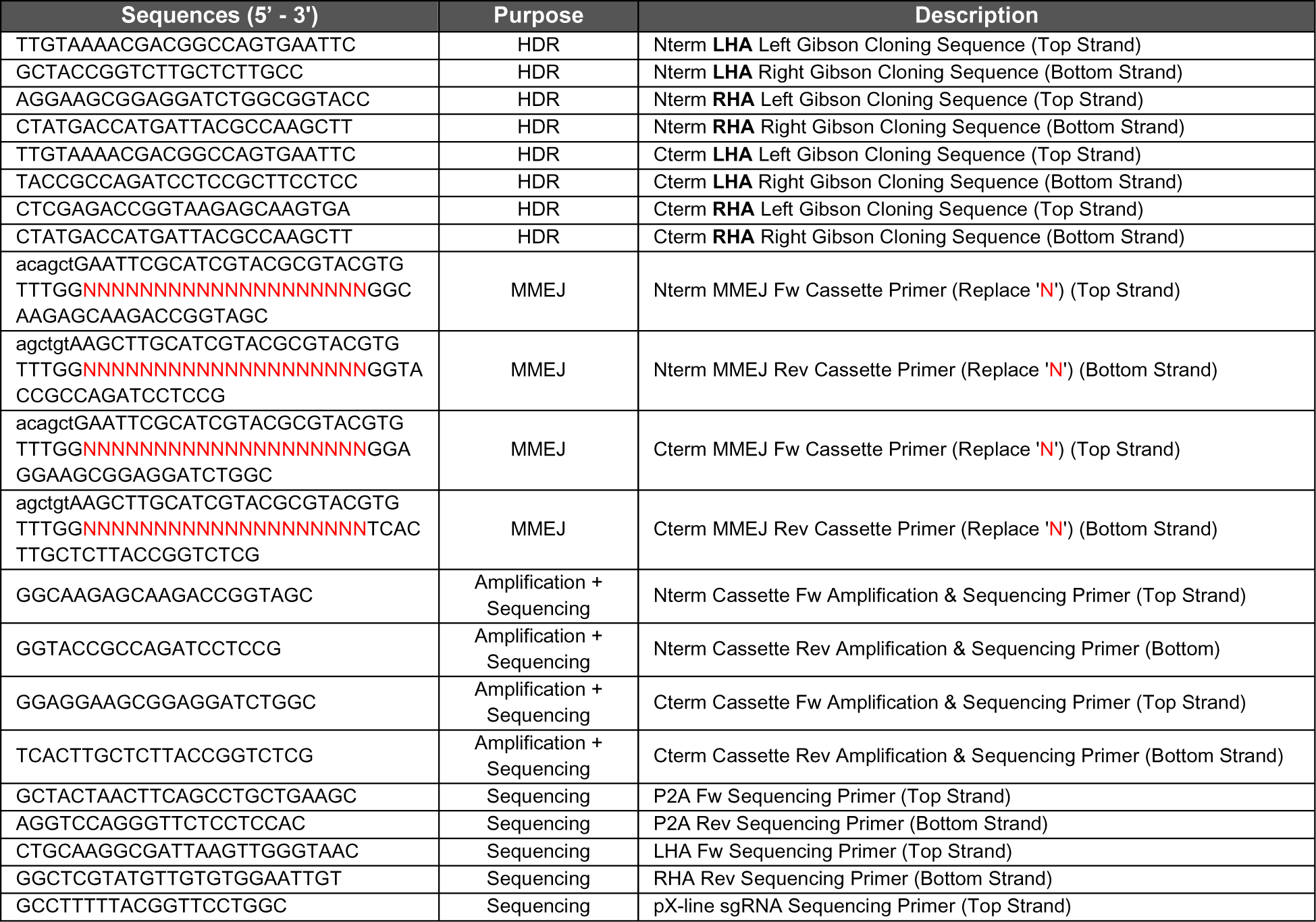
qTAG relevant sequences and primers.

For MMEJ-based qTAG repair constructs, flanking homology of 10-20 bp around the sgRNA cut site was directly added to the cassette amplification primers used to amplify the preferred qTAG cassette. PITCh sgRNA sequences were also included in the primers following the homology sequences to allow for cassette release upon cleavage. The qTAG MMEJ primer design scaffold is outlined in Table 2. Following the PCR process with the MMEJ primers and the selected qTAG cassette, the resultant cassette fragment underwent digestion and was subsequently restriction-cloned into a pUC19 backbone.

In both the HDR and MMEJ cases, the repair constructs were purified (Plasmid Plus Midi Kit, 12945, QIAGEN) and sequence verified via Sanger sequencing using the sequencing primers outlined in Table 2 prior to introduction into cells. Source gene designs of all of the genes targeted in this paper are highlighted in Table 3 with their corresponding qTAG repair variants summarized in Table 4. Additionally, supplementary example annotated sequence data files are provided for a more detailed view of knock-in design and the resultant qTAG repair constructs (Supp. Data 1). For detailed, step-by-step instructions on gene design, plasmid construction, cell culture, and the validation of knock-ins using these qTAG cassettes, please refer to https://www.biorxiv.org/content/10.1101/2023.11.01.565029v2.

**Table 3.**
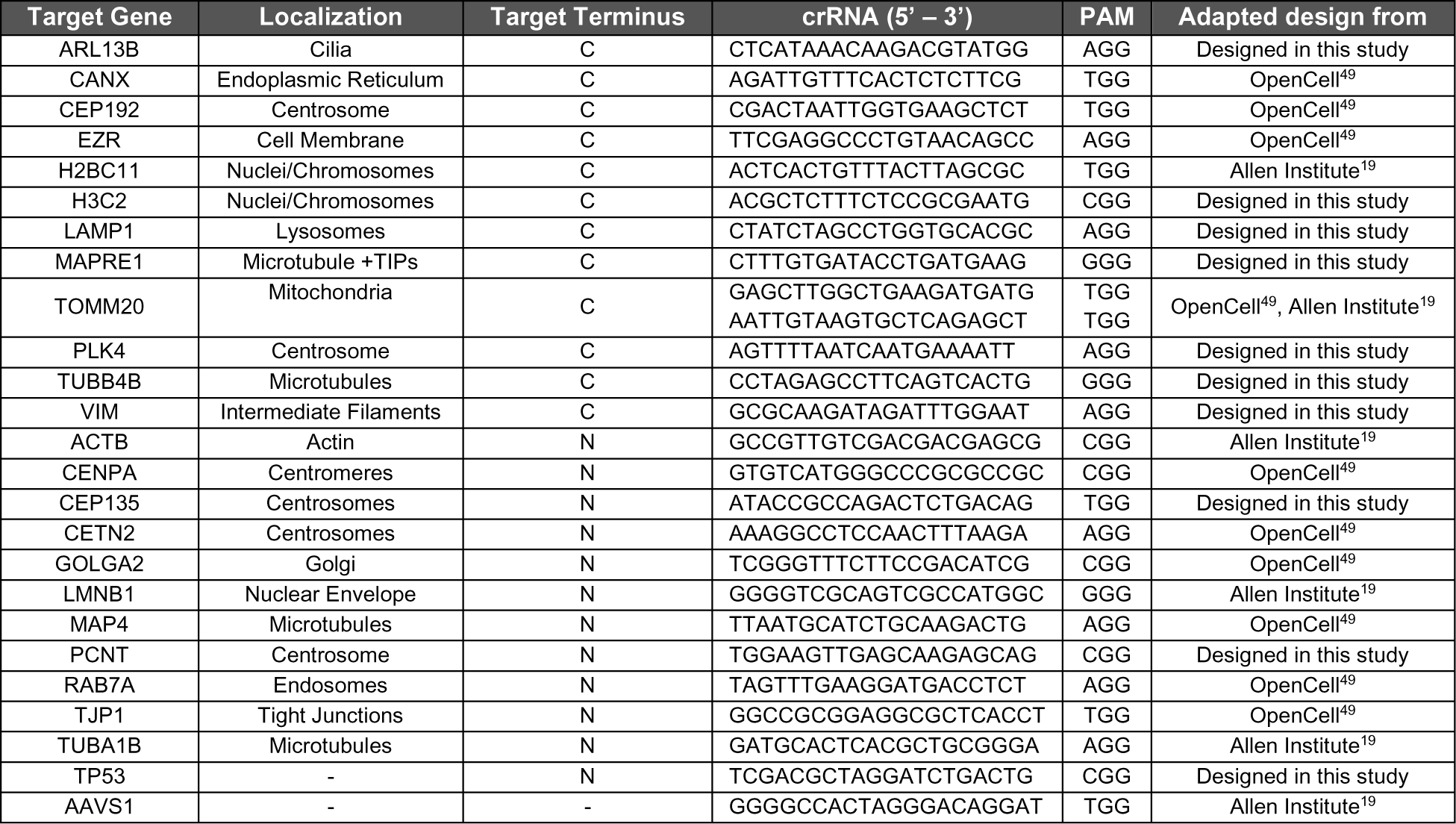
Gene Design Information.

**Table 4.** Plasmids for the qTAG system.

| Plasmid Name | Terminus | Tag | Marker | Plasmid Information and Map |
| --- | --- | --- | --- | --- |
| qTAG-N-Blast-mNeon | N | mNeonGreen | Blasticidin | https://www.addgene.org/207676/ |
| qTAG-N-Blast-moxGFP | N | moxGFP | Blasticidin | https://www.addgene.org/207677/ |
| qTAG-N-Blast-mScarlet | N | mScarlet | Blasticidin | https://www.addgene.org/207678/ |
| qTAG-N-Blast-sTagRFP | N | super-TagRFP | Blasticidin | https://www.addgene.org/207679/ |
| qTAG-N-Blast-miRFP670nano3 | N | miRFP670nano3 | Blasticidin | https://www.addgene.org/207680/ |
| qTAG-N-Blast-miniTurbo | N | miniTurbo | Blasticidin | https://www.addgene.org/207681/ |
| qTAG-N-Blast-ultraID | N | ultraID | Blasticidin | https://www.addgene.org/207682/ |
| qTAG-N-Blast-dTAG | N | dTAG | Blasticidin | https://www.addgene.org/207683/ |
| qTAG-N-Blast-3xFLAG | N | 3xFLAG | Blasticidin | https://www.addgene.org/207684/ |
| qTAG-N-Blast-3xHA | N | 3xHA | Blasticidin | https://www.addgene.org/207685/ |
| qTAG-N-Blast-V5 | N | V5 | Blasticidin | https://www.addgene.org/207686/ |
| qTAG-N-Puro-mNeon | N | mNeonGreen | Puromycin | https://www.addgene.org/207687/ |
| qTAG-N-Puro-moxGFP | N | moxGFP | Puromycin | https://www.addgene.org/207688/ |
| qTAG-N-Puro-mScarlet | N | mScarlet | Puromycin | https://www.addgene.org/207689/ |
| qTAG-N-Puro-sTagRFP | N | super-TagRFP | Puromycin | https://www.addgene.org/207690/ |
| qTAG-N-Puro-miRFP670nano3 | N | miRFP670nano3 | Puromycin | https://www.addgene.org/207691/ |
| qTAG-N-Puro-miniTurbo | N | miniTurbo | Puromycin | https://www.addgene.org/207692/ |
| qTAG-N-Puro-ultraID | N | ultraID | Puromycin | https://www.addgene.org/207693/ |
| qTAG-N-Puro-dTAG | N | dTAG | Puromycin | https://www.addgene.org/207694/ |
| qTAG-N-Puro-3xFLAG | N | 3xFLAG | Puromycin | https://www.addgene.org/207695/ |
| qTAG-N-Puro-3xHA | N | 3xHA | Puromycin | https://www.addgene.org/207696/ |
| qTAG-N-Puro-V5 | N | V5 | Puromycin | https://www.addgene.org/207697/ |
| qTAG-N-Zeo-mNeon | N | mNeonGreen | Zeocin | https://www.addgene.org/207698/ |
| qTAG-N-Zeo-moxGFP | N | moxGFP | Zeocin | https://www.addgene.org/207699/ |
| qTAG-N-Zeo-mScarlet | N | mScarlet | Zeocin | https://www.addgene.org/207700/ |
| qTAG-N-Zeo-sTagRFP | N | super-TagRFP | Zeocin | https://www.addgene.org/207701/ |
| qTAG-N-Zeo-miRFP670nano3 | N | miRFP670nano3 | Zeocin | https://www.addgene.org/207702/ |
| qTAG-N-Zeo-miniTurbo | N | miniTurbo | Zeocin | https://www.addgene.org/207703/ |
| qTAG-N-Zeo-ultraID | N | ultraID | Zeocin | https://www.addgene.org/207704/ |
| qTAG-N-Zeo-dTAG | N | dTAG | Zeocin | https://www.addgene.org/207705/ |
| qTAG-N-Zeo-3xFLAG | N | 3xFLAG | Zeocin | https://www.addgene.org/207706/ |
| qTAG-N-Zeo-3xHA | N | 3xHA | Zeocin | https://www.addgene.org/207707/ |
| qTAG-N-Zeo-V5 | N | V5 | Zeocin | https://www.addgene.org/207708/ |
| qTAG-C-mNeon-Blast | C | mNeonGreen | Blasticidin | https://www.addgene.org/207709/ |
| qTAG-C-moxGFP-Blast | C | moxGFP | Blasticidin | https://www.addgene.org/207710/ |
| qTAG-C-mScarlet-Blast | C | mScarlet | Blasticidin | https://www.addgene.org/207711/ |
| qTAG-C-sTagRFP-Blast | C | super-TagRFP | Blasticidin | https://www.addgene.org/207712/ |
| qTAG-C-miRFP670nano3-Blast | C | miRFP670nano3 | Blasticidin | https://www.addgene.org/207713/ |
| qTAG-C-miniTurbo-Blast | C | miniTurbo | Blasticidin | https://www.addgene.org/207714/ |
| qTAG-C-ultraID-Blast | C | ultraID | Blasticidin | https://www.addgene.org/207715/ |
| qTAG-C-dTAG-Blast | C | dTAG | Blasticidin | https://www.addgene.org/207716/ |
| qTAG-C-3xFLAG-Blast | C | 3xFLAG | Blasticidin | https://www.addgene.org/207717/ |
| qTAG-C-3xHA-Blast | C | 3xHA | Blasticidin | https://www.addgene.org/207718/ |
| qTAG-C-V5-Blast | C | V5 | Blasticidin | https://www.addgene.org/207719/ |
| qTAG-C-mNeon-Puro | C | mNeonGreen | Puromycin | https://www.addgene.org/207720/ |
| qTAG-C-moxGFP-Puro | C | moxGFP | Puromycin | https://www.addgene.org/207721/ |
| qTAG-C-mScarlet-Puro | C | mScarlet | Puromycin | https://www.addgene.org/207722/ |
| qTAG-C-sTagRFP-Puro | C | super-TagRFP | Puromycin | https://www.addgene.org/207723/ |
| qTAG-C-miRFP670nano3-Puro | C | miRFP670nano3 | Puromycin | https://www.addgene.org/207724/ |
| qTAG-C-miniTurbo-Puro | C | miniTurbo | Puromycin | https://www.addgene.org/207725/ |
| qTAG-C-ultraID-Puro | C | ultraID | Puromycin | https://www.addgene.org/207726/ |
| qTAG-C-dTAG-Puro | C | dTAG | Puromycin | https://www.addgene.org/207727/ |
| qTAG-C-3xFLAG-Puro | C | 3xFLAG | Puromycin | https://www.addgene.org/207728/ |
| qTAG-C-3xHA-Puro | C | 3xHA | Puromycin | https://www.addgene.org/207729/ |
| qTAG-C-V5-Puro | C | V5 | Puromycin | https://www.addgene.org/207730/ |
| qTAG-C-mNeon-Zeo | C | mNeonGreen | Zeocin | https://www.addgene.org/207731/ |
| qTAG-C-moxGFP-Zeo | C | moxGFP | Zeocin | https://www.addgene.org/207732/ |
| qTAG-C-mScarlet-Zeo | C | mScarlet | Zeocin | https://www.addgene.org/207733/ |
| qTAG-C-sTagRFP-Zeo | C | super-TagRFP | Zeocin | https://www.addgene.org/207734/ |
| qTAG-C-miRFP670nano3-Zeo | C | miRFP670nano3 | Zeocin | https://www.addgene.org/207735/ |
| qTAG-C-miniTurbo-Zeo | C | miniTurbo | Zeocin | https://www.addgene.org/207736/ |
| qTAG-C-ultraID-Zeo | C | ultraID | Zeocin | https://www.addgene.org/207737/ |
| qTAG-C-dTAG-Zeo | C | dTAG | Zeocin | https://www.addgene.org/207738/ |
| qTAG-C-3xFLAG-Zeo | C | 3xFLAG | Zeocin | https://www.addgene.org/207739/ |
| qTAG-C-3xHA-Zeo | C | 3xHA | Zeocin | https://www.addgene.org/207740/ |
| qTAG-C-V5-Zeo | C | V5 | Zeocin | https://www.addgene.org/207741/ |
| pEF1α-Cre-PGK-Puro | - | Cre Recombinase | Puromycin | https://www.addgene.org/207742/ |
| pEF1α-Cre-2A-Puro | - | Cre Recombinase | Puromycin | https://www.addgene.org/207743/ |
| pEF1α-Cre-2A-moxGFP | - | Cre Recombinase | moxGFP | https://www.addgene.org/207744/ |
| pEF1α-Cre-2A-mScarlet | - | Cre Recombinase | mScarlet | https://www.addgene.org/207745/ |
| pEF1α-Cre-2A-mTagBFP2 | - | Cre Recombinase | mTagBFP2 | https://www.addgene.org/207746/ |
| pEF1α-Cre-2A-miRFP670nano3 | - | Cre Recombinase | miRFP670nano3 | https://www.addgene.org/207747/ |
| pX330 | - | Cas9 + sgRNA | - | https://www.addgene.org/42230/ |
| pX330-PITCH | - | Cas9 + sgRNA + PITCH sgRNA | - | https://www.addgene.org/127875/ |
| pX330-PITCh-ACTB | - | Cas9 + sgRNA + PITCh sgRNA | - | https://www.addgene.org/207748/ |
| qTAG-N-Solo-mNeon-ACTB | N | mNeonGreen | - | https://www.addgene.org/207749/ |
| qTAG-N-Solo-mScarlet-ACTB | N | mScarlet | - | https://www.addgene.org/207750/ |
| qTAG-N-Blast-mNeon-ACTB | N | mNeonGreen | Blasticidin | https://www.addgene.org/207751/ |
| qTAG-N-Puro-mNeon-ACTB | N | mNeonGreen | Puromycin | https://www.addgene.org/207752/ |
| qTAG-N-Blast-mScarlet-ACTB | N | mScarlet | Blasticidin | https://www.addgene.org/207753/ |
| qTAG-N-Puro-mScarlet-ACTB | N | mScarlet | Puromycin | https://www.addgene.org/207754/ |
| pX330-PITCh-H2BC11 | - | Cas9 + sgRNA + PITCh sgRNA | - | https://www.addgene.org/207755/ |
| qTAG-C-Solo-mScarlet-H2BC11 | C | mScarlet | - | https://www.addgene.org/207756/ |
| qTAG-C-moxGFP-Blast-H2BC11 | C | moxGFP | Blasticidin | https://www.addgene.org/207757/ |
| qTAG-C-moxGFP-Puro-H2BC11 | C | moxGFP | Puromycin | https://www.addgene.org/207758/ |
| qTAG-C-moxGFP-Zeo-H2BC11 | C | moxGFP | Zeocin | https://www.addgene.org/207759/ |
| qTAG-C-moxGFP-Puro-H2BC11-MMEJ | C | moxGFP | Puromycin | https://www.addgene.org/207760/ |
| qTAG-C-mScarlet-Puro-H2BC11 | C | mScarlet | Puromycin | https://www.addgene.org/207761/ |
| qTAG-C-miRFP670nano3-Blast-H2BC11 | C | miRFP670nano3 | Blasticidin | https://www.addgene.org/207762/ |
| pX330-PITCh-TUBA1B | - | Cas9 + sgRNA + PITCh sgRNA | - | https://www.addgene.org/207763/ |
| qTAG-N-Solo-mScarlet-TUBA1B | N | mScarlet | - | https://www.addgene.org/207764/ |
| qTAG-N-Solo-sTagRFP-TUBA1B | N | super-TagRFP | - | https://www.addgene.org/207765/ |
| qTAG-N-Blast-moxGFP-TUBA1B | N | moxGFP | Blasticidin | https://www.addgene.org/207766/ |
| qTAG-N-Puro-moxGFP-TUBA1B | N | moxGFP | Puromycin | https://www.addgene.org/207767/ |
| qTAG-N-Blast-sTagRFP-TUBA1B | N | super-TagRFP | Blasticidin | https://www.addgene.org/207768/ |
| qTAG-N-Puro-sTagRFP-TUBA1B | N | super-TagRFP | Puromycin | https://www.addgene.org/207769/ |
| pX330-PITCh-LMNB1 | - | Cas9 + sgRNA + PITCh sgRNA | - | https://www.addgene.org/207770/ |
| qTAG-N-Solo-mScarlet-LMNB1 | N | mScarlet | - | https://www.addgene.org/207771/ |
| qTAG-N-Blast-moxGFP-LMNB1 | N | moxGFP | Blasticidin | https://www.addgene.org/207772/ |
| qTAG-N-Puro-moxGFP-LMNB1 | N | moxGFP | Puromycin | https://www.addgene.org/207773/ |
| qTAG-N-Blast-miniTurbo-LMNB1 | N | miniTurbo | Blasticidin | https://www.addgene.org/207774/ |
| qTAG-N-Blast-ultraID-LMNB1 | N | ultraID | Blasticidin | https://www.addgene.org/207775/ |
| qTAG-N-Blast-dTAG-LMNB1 | N | dTAG | Blasticidin | https://www.addgene.org/207776/ |
| qTAG-N-Blast-3xFLAG-LMNB1 | N | 3xFLAG | Blasticidin | https://www.addgene.org/207777/ |
| qTAG-N-Blast-3xHA-LMNB1 | N | 3xHA | Blasticidin | https://www.addgene.org/207778/ |
| qTAG-N-Blast-V5-LMNB1 | N | V5 | Blasticidin | https://www.addgene.org/207779/ |
| pX330-PITCh-H3C2 | - | Cas9 + sgRNA + PITCh sgRNA | - | https://www.addgene.org/207780/ |
| qTAG-C-Solo-mScarlet-H3C2 | C | mScarlet | - | https://www.addgene.org/207781/ |
| qTAG-C-mNeon-Blast-H3C2 | C | mNeonGreen | Blasticidin | https://www.addgene.org/207782/ |
| qTAG-C-moxGFP-Puro-H3C2 | C | moxGFP | Puromycin | https://www.addgene.org/207783/ |
| qTAG-C-miRFP670nano3-Blast-H3C2 | C | miRFP670nano3 | Blasticidin | https://www.addgene.org/207784/ |
| pX330-PITCh-TUBB4B | - | Cas9 + sgRNA + PITCh sgRNA | - | https://www.addgene.org/207785/ |
| qTAG-C-mNeon-Blast-TUBB4B | C | mNeonGreen | Blasticidin | https://www.addgene.org/207786/ |
| pX330-LAMP1 | - | Cas9 + sgRNA + PITCh sgRNA | - | https://www.addgene.org/207787/ |
| qTAG-C-moxGFP-Puro-LAMP1 | C | moxGFP | Puromycin | https://www.addgene.org/207788/ |
| pX330-PITCh-TOMM20 | - | Cas9 + sgRNA + PITCh sgRNA | - | https://www.addgene.org/207789/ |
| qTAG-C-mNeon-Puro-TOMM20 | C | mNeonGreen | Puromycin | https://www.addgene.org/207790/ |
| pX330-PITCh-GOLGA2 | - | Cas9 + sgRNA + PITCh sgRNA | - | https://www.addgene.org/207791/ |
| qTAG-N-Puro-moxGFP-GOLGA2 | N | moxGFP | Puromycin | https://www.addgene.org/207792/ |
| pX330-PITCh-MAPRE1 | - | Cas9 + sgRNA + PITCh sgRNA | - | https://www.addgene.org/207793/ |
| qTAG-C-moxGFP-Puro-MAPRE1 | C | moxGFP | Puromycin | https://www.addgene.org/207794/ |
| pX330-PITCh-TOMM20-2 | C | Cas9 + sgRNA + PITCh sgRNA | - | Being submitted to Addgene |
| qTAG-N-Solo-mStayGold | N | mStayGold | - | Being submitted to Addgene |
| qTAG-N-Puro-mStayGold | N | mStayGold | Puromycin | Being submitted to Addgene |
| qTAG-N-Blast-mStayGold | N | mStayGold | Blasticidin | Being submitted to Addgene |
| qTAG-N-Zeo-mStayGold | N | mStayGold | Zeocin | Being submitted to Addgene |
| qTAG-C-Solo-mStayGold | C | mStayGold | - | Being submitted to Addgene |
| qTAG-C-mStayGold-Puro | C | mStayGold | Puromycin | Being submitted to Addgene |
| qTAG-C-mStayGold-Blast | C | mStayGold | Blasticidin | Being submitted to Addgene |
| qTAG-C-mStayGold-Zeo | C | mStayGold | Zeocin | Being submitted to Addgene |
| qTAG-C-mStayGold-EFs-Puro | C | mStayGold | Puromycin | Being submitted to Addgene |
| qTAG-C-mStayGold-EFs-Blast | C | mStayGold | Blasticidin | Being submitted to Addgene |
| qTAG-C-mStayGold-EFs-Zeo | C | mStayGold | Zeocin | Being submitted to Addgene |
| qTAG-C-mScarlet-EFs-Puro | C | mScarlet | Puromycin | Being submitted to Addgene |
| qTAG-C-mScarlet-EFs-Blast | C | mScarlet | Blasticidin | Being submitted to Addgene |
| qTAG-C-mScarlet-EFs-Zeo | C | mScarlet | Zeocin | Being submitted to Addgene |
| qTAG-KO-Puro | N | - | Puromycin | Being submitted to Addgene |
| qTAG-KO-Blast | N | - | Blasticidin | Being submitted to Addgene |
| qTAG-KO-PGK-Puro | N | - | Puromycin | Being submitted to Addgene |
| qTAG-KO-PGK-Blast | N | - | Blasticidin | Being submitted to Addgene |
| qTAG-AAVS1-Puro-Ef1a | - | - | Puromycin | Being submitted to Addgene |
| qTAG-AAVS1-Blast-Ef1a | - | - | Blasticidin | Being submitted to Addgene |
| qTAG-AAVS1-Zeo-Ef1a | - | - | Zeocin | Being submitted to Addgene |
| qTAG-AAVS1-Puro-PGK | - | - | Puromycin | Being submitted to Addgene |
| qTAG-AAVS1-Blast-PGK | - | - | Blasticidin | Being submitted to Addgene |
| qTAG-AAVS1-Zeo-PGK | - | - | Zeocin | Being submitted to Addgene |
| qTAG-AAVS1-Puro-TetO | - | - | Puromycin | Being submitted to Addgene |
| qTAG-AAVS1-Blast-TetO | - | - | Blasticidin | Being submitted to Addgene |
| qTAG-AAVS1-Zeo-TetO | - | - | Zeocin | Being submitted to Addgene |
| pX330-AAVS1 | - | Cas9 + sgRNA | - | Being submitted to Addgene |
| pX330-PITCh-ARL13B | - | Cas9 + sgRNA + PITCh sgRNA | - | Being submitted to Addgene |
| pX330-CANX | - | Cas9 + sgRNA | - | Being submitted to Addgene |
| pX330-CENPA | - | Cas9 + sgRNA | - | Being submitted to Addgene |
| pX330-PCNT | - | Cas9 + sgRNA | - | Being submitted to Addgene |
| pX330-CEP135 | - | Cas9 + sgRNA | - | Being submitted to Addgene |
| pX330-CEP192 | - | Cas9 + sgRNA | - | Being submitted to Addgene |
| pX330-CETN2 | - | Cas9 + sgRNA | - | Being submitted to Addgene |
| pX330-EZR | - | Cas9 + sgRNA | - | Being submitted to Addgene |
| pX330-PITCh-MAP4 | - | Cas9 + sgRNA + PITCh sgRNA | - | Being submitted to Addgene |
| pX330-RAB7A | - | Cas9 + sgRNA | - | Being submitted to Addgene |
| pX330-TJP1 | - | Cas9 + sgRNA | - | Being submitted to Addgene |
| pX330-PITCh-VIM | - | Cas9 + sgRNA + PITCh sgRNA | - | Being submitted to Addgene |
| pX330-TP53 | - | Cas9 + sgRNA | - | Being submitted to Addgene |
| qTAG-AAVS1-Puro-Ef1a-moxGFP | - | moxGFP | Puromycin | Being submitted to Addgene |
| qTAG-AAVS1-Puro-Ef1a-mScarlet | - | mScarlet | Puromycin | Being submitted to Addgene |
| qTAG-KO-Puro-TP53 | N | - | Puromycin | Being submitted to Addgene |
| qTAG-C-mStayGold-EFs-Puro-PLK4 | C | mStayGold | Puromycin | Being submitted to Addgene |
| qTAG-C-mStayGold-EFs-Blast-ARL13B | C | mStayGold | Blasticidin | Being submitted to Addgene |
| qTAG-C-Solo-miRFP670nano3-H2BC11 | C | miRFP670nano3 | - | Being submitted to Addgene |
| qTAG-C-Solo-miRFP670nano3-H3C2 | C | miRFP670nano3 | - | Being submitted to Addgene |
| qTAG-N-Solo-mStayGold-ACTB | N | mStayGold | - | Being submitted to Addgene |
| qTAG-N-Solo-mStayGold-TUBA1B | N | mStayGold | - | Being submitted to Addgene |
| qTAG-C-Solo-mStayGold-H2BC11 | C | mStayGold | - | Being submitted to Addgene |
| qTAG-C-Solo-mStayGold-H3C2 | C | mStayGold | - | Being submitted to Addgene |
| qTAG-C-Solo-mStayGold-EZR | C | mStayGold | - | Being submitted to Addgene |
| qTAG-N-Solo-mScarlet-MAP4 | N | mScarlet | - | Being submitted to Addgene |
| qTAG-N-Solo-mStayGold-MAP4 | N | mStayGold | - | Being submitted to Addgene |
| qTAG-C-Solo-mStayGold-MAPRE1 | C | mStayGold | - | Being submitted to Addgene |
| qTAG-C-Solo-mScarlet-MAPRE1 | C | mScarlet | - | Being submitted to Addgene |
| qTAG-N-Solo-mStayGold-CENPA | N | mStayGold | - | Being submitted to Addgene |
| qTAG-N-Solo-mStayGold-GOLGA2 | N | mStayGold | - | Being submitted to Addgene |
| qTAG-C-Solo-mStayGold-CANX | C | mStayGold | - | Being submitted to Addgene |
| qTAG-C-Solo-mStayGold-TOMM20 | C | mStayGold | - | Being submitted to Addgene |
| qTAG-N-Solo-mStayGold-TJP1 | N | mStayGold | - | Being submitted to Addgene |
| qTAG-N-Solo-mStayGold-RAB7A | N | mStayGold | - | Being submitted to Addgene |
| qTAG-C-Solo-mStayGold-VIM | C | mStayGold | - | Being submitted to Addgene |
| qTAG-N-Solo-mStayGold-LMNB1 | N | mStayGold | - | Being submitted to Addgene |
| qTAG-N-Solo-mStayGold-CEP135 | N | mStayGold | - | Being submitted to Addgene |
| qTAG-N-Solo-mStayGold-PCNT | N | mStayGold | - | Being submitted to Addgene |
| qTAG-N-Solo-mStayGold-CETN2 | N | mStayGold | - | Being submitted to Addgene |
| qTAG-C-Solo-mStayGold-CEP192 | C | mStayGold | - | Being submitted to Addgene |
| qTAG-C-Solo-mStayGold-LAMP1 | C | mStayGold | - | Being submitted to Addgene |
| qTAG-C-Solo-mStayGold-ARL13B | C | mStayGold | - | Being submitted to Addgene |

### Cell culture

The human cell lines used in this study included: HAP1 cells (Horizon Genomics, cat. No. C631, RRID: CVCL_Y019), HEK293T cells (ATCC, cat. No. CRL-3216, RRID: CVCL_0063), ARPE-19 cells (ATCC, cat. No. CRL-3216, RRID: CVCL_0145), RPE-1 cells (ATCC, cat. No. CRL-4000, RRID: CVCL_4388), U-2 OS cells (ATCC, cat. No. HTB-96, RRID: CVCL_0042), H9 cells (WiCell, cat. WA09, RRID: CVCL_9773). RPE-1 with a knocked-out PAC (puromycin N-acetyltransferase) gene was obtained as a gift from Dr. Daniel Durocher’s laboratory. DMEM (11965084, Gibco) supplemented with 10% FBS was used to culture HAP1 and HEK293T and cells. For U-2 OS cells, McCoy’s 5A medium (16600082, Gibco) supplemented with 10% FBS was used. DMEM/F-12 (11320033, Gibco) supplemented with 10% FBS was used to culture ARPE-19 and RPE-1 PAC (-) cells. mTeSR1 (100-0276, StemCell Tech) and CloneR (05889, StemCell Tech) were used for regular and clonal culture H9 cells, respectively. All cells were grown in a tissue culture incubator with humidity at 37 °C and 5% CO2.

### Transfection and editing

Chemical delivery of editing components via Lipofectamine 3000 into tissue culture cell lines began with seeding approximately 500,000 cells (or enough to reach 70-80% confluency the next day) per well in a 6-well plate. Each experiment included a well designated for transfection and selection controls, positioned alongside the well containing the gene-tagging target. The subsequent day, co-transfection of pX-CRISPR and qTAG repair plasmids was performed at a 1:1 ratio using 4 μg of total DNA (2 μg cut + 2 μg repair) in Opti-MEM Reduced Serum Medium (31985070, Gibco) with Lipofectamine 3000 (L3000008, Invitrogen). Media replacement occurred the next day and an editing period of 72 h post-transfection was allowed prior to selection enrichment. It was crucial to transfer cells to a larger vessel (6-well > 10cm dish) if confluency was reached to ensure proper cell growth and prevent contact inhibition.

The delivery of editing components via Neon electroporation (MPK10096, MPK5000, ThermoFisher) and RNPs into H9 human stem cells involved several steps. Initially, H9 cells were dissociated into single cells using TrypLE (12605028, ThermoFisher) and then collected via centrifugation and resuspended in mTeSR1. RNP complexes were assembled on ice by incubating purified Cas9 protein with the sgRNA, followed by addition to cells at a cell concentration of 1 x 10^7^ cells/mL in 10 μL. Electroporation was carried out using specific parameters (1100 V, 30 ms, 1 Pulse) using 10 μL Neon tips, and cells were rapidly transferred to media containing CloneR in a Matrigel-coated well. The following day, media replacement was performed, and an editing period of up to 72 h was allowed before selection enrichment.

### Enrichment of edits via mammalian antibiotic selection or FACS

After editing, the culture media was replaced with media supplemented with the mammalian antibiotic corresponding to the tagging cassette used. A range of working concentrations was tested for ARPE-19, RPE-1, HEK293T, HAP1, U-2 OS, and H9 cells. To enrich for knocked-in cells in human tissue culture cell lines, concentrations of 5-10 μg/mL Blasticidin, 0.5-1 μg/mL Puromycin, and 100-400 μg/mL Zeocin were used, with complete selection being observed in 4-7 days, 2-3 days, and 10-15 days, respectively. For H9 stem cells, concentrations of 3 μg/mL Blasticidin, 0.5 μg/mL Puromycin, and 25-50 μg/mL Zeocin were used, with complete selection being observed in 7-8 days, 2-3 days, and 10-15 days, respectively. Once all untransfected control cells had been eradicated, the resultant resistant cells represent the initial enriched tagged pool. It was critical to note that the initial number of remaining cells in some cases were limited, depending on factors such as guide or transfection efficiency. In such cases (Certain gene designs including PLK4, ARL13B, TOMM20), an additional 2-5 days were allotted for growth of potential resistant colonies. If it wasn’t feasible to incorporate mammalian selectable markers or if a Tag-Only approach was employed, as seen in the mStayGold experiments, edited cell enrichment was alternatively performed using FACS (BD FACS Aria Fusion). Fluorescence thresholds and gates were based on the negative-control, unedited population.

### Cre selectable marker deletion

Cre selectable marker excision was carried out after the generation of a selected pool of edited cells. 500,000 of the target tagged cell line was seeded per well in a 6-well plate to achieve ∼80% confluency the following day. Cre recombinase delivery involved transfection of the chosen Cre encoding plasmid, Cre-Puro in the case of Fig. 3, using Lipofectamine 3000 as previously described. The following day, the selection of cells commenced either through fluorescence or puromycin resistance, dependent on the Cre plasmid used, to enrich for cells expressing the Cre recombinase, resulting in the generation of a tagged pool of cells with the integrated selectable marker being removed. Precaution was taken to remove the selection immediately after the elimination of the control cells to avoid enriching for permanent integrations of the Cre plasmid itself.

### Clonal cell line generation

To isolate clones, ∼350-500 cells from the selected pool of edited cells were seeded into a 15 cm dish. 7 - 10 days were allotted for the formation of distinct colonies. Cellular clones were then isolated using cloning cylinders (CLS31666, MiliporeSigma) coated in sterile silicone grease (85410, MiliporeSigma), where the clones could be isolated, trypsinized, and transferred for further propagation.

### Flow Cytometry

Flow analysis of the edited fluorescent populations was carried out to assess tagging efficiency and enrichment using the qTAG cassettes. Control, initial tagged pools, and selected pools of cells were resuspended in a PBS, 2% FBS, 0.5 mM EDTA buffer and filtered through a cell strainer (352235, Corning) to make a single cell suspension. Samples were then subjected to analysis on an Attune NxT flow cytometer (ThermoFisher) where size and fluorescence thresholds and gates were based on the negative-control, unedited population. For each sample, ∼200, 000 events were recorded in a technical triplicate across 3 biological replicates.

### Selection Assay

For selection assay experiments, 200,000 cells previously transfected with editing reagents were seeded in 6-well plates. The next day medium was removed and medium containing the indicated selection was added. The media was refreshed every 3–4 days to ensure continued selection. After 7 days, plates were rinsed once with PBS and fixed and stained with 0.5% crystal violet (46364, MilliporeSigma) in 20% methanol for at least 20 min. Plates were washed extensively with water, dried, and scanned.

### Allelic screening with genomic junction PCR

To check for the allelic integration status of the cassettes, genomic DNA was extracted from 10,000 – 50,000 target cells by resuspending them in a QuickExtract DNA Extraction Solution (LGN-QE09050, Lucigen). PCR amplification of the targeted junction from the wild-type and tagged cells was performed using primers designed to anneal outside of the homology arms of the specific gene. The resulting PCR products were analyzed on an agarose gel, where it was expected that the size of the amplicon from the tagged alleles would be proportional to the qTAG insertion cassette size plus the wild-type amplicon size. For further validation of framing and the absence of mutations, the amplicons were gel purified using the QIAquick Gel Extraction Kit (28706, QIAGEN) and subjected to sanger sequencing.

### Immunoblot Analyses

For immunoblot analyses, cells were seeded into 6-well plates and cultured until confluency was reached. The indicated cells were then scraped and lysed on ice for 10 min with RIPA buffer (150 mM NaCl, 1% Triton X-100, 0.5% sodium deoxycolate, 0.1% SDS, 50 mM Tris-HCl at pH 8.0, 10 mM NaF, 1 mM Na3VO4, 1 mM EDTA and 1 mM EGTA) containing a protease inhibitor cocktail (Sigma-Aldrich, P8340) and centrifuged at 12,000 r.p.m. for 10 min at 4 °C. To quantitate the protein, a fraction of the supernatant was subjected to a Bradford assay (Bradford Protein Assay Kit, 23200, ThermoFisher). The rest of the supernatant was mixed with 4x SDS sample buffer (250 mM Tris-HCl at pH 6.8, 8% SDS, 40% glycerol and 0.04% bromophenol blue) and 10 mM DTT (Amresco, 0281-25G). The mixtures were boiled for 5 min. To ascertain the presence of the endogenous protein or its endogenous tag, 10 ug of protein of each sample was loaded in 10% SDS–polyacrylamide gel, electrophoresed, and transferred to PVDF. The total protein was detected by staining with PonceauS (P7170, MilliporeSigma) prior to scanning. The membranes were blocked with a blocking solution (5% nonfat milk in 0.1% Tween-20 in TBS) for 1 h, incubated with primary antibodies diluted in blocking solution overnight at 4 °C, washed four times x 5min with 0.1% Tween-20 in TBS (TBST), incubated with fluorescence-based secondary antibodies (LI-COR Biosciences) in blocking solution for 1 h, then washed again four times in TBST. The membrane was then dried for 1 h at room temperature prior to being imaged using an Odyssey CLx imager (LI-COR Biosciences). Antibodies used for immunoblotting in this study included: Streptavidin (926-68079, LI-COR Biosciences), LMNB1 (AB16048, Abcam), FLAG (14793, Cell Signaling Technology), HA (901501, BioLegend), V5 (R960-25, ThermoFisher), TP53 (sc-126, Santa Cruz), and GAPDH (G9545-100UL, MilliporeSigma).

### Fluorescence staining, fixed imaging, and live-cell Imaging

For fixed cell imaging, cells were grown as indicated on No. 1.5 coverslips, washed once with PBS, and fixed with either 4% PFA or –20 °C methanol for 10 min. The subsequent steps were performed at room temperature unless otherwise denoted. The coverslips were then rinsed with 0.1% Triton X-100 in PBS (PBST) 3x, then permeabilized (in the case of PFA fixation) with 0.3% PBST for 20 min. 3x more washes in 0.1% PBST were carried out prior to blocking with 5% BSA in PBST for 30 min. If the experiment called for primary antibodies, samples were incubated with primary antibodies diluted in blocking solution for 1 h, then washed 3x with 0.1% PBST. Complementary secondary antibodies (Alexa Secondaries, ThermoFisher), combined with any cellular stains such as DAPI, were incubated 1 h. Coverslips were washed 3x 0.1% PBST and mounted on slides using Prolong Gold (P36930, ThermoFisher). Antibodies used for immunofluorescence in this study included: Previously listed antibodies, ARL13B (sc-515784, Santa Cruz), CEP135 (produced in house). For the selectable knock-out experiments, Nutlin-3a (18585, Cayman Chemical) was used at 10 μM to induce TP53 accumulation in cells for detection.

For fixed imaging, spinning-disk confocal microscopy was performed using either a Nikon W1-CSU-SoRa or a Nikon CrestOptics-X-Light-DeepSIM microscope equipped with a CFI Plan Apo IR 60XC WI water immersion objective (NA 1.27). Each field was acquired where the whole cell volume of 20 μm could be captured with a z-step of 0.3 μm. Displayed are maximum intensity projections. For live imaging, 100,000 cells were seeded per well in a four-well Lab-Tek II chamber slide. To capture live-cell snapshots, the cells were treated with SiR live cell dyes targeting actin, tubulin, or DNA (Spirochrome) at a concentration of 100 nM for 2 h before imaging. Cells were maintained at 37°C and 5% CO2 using a Tokai Hit environmental control chamber. For the photobleaching assay, RPE-1 PAC (-) cells endogenously tagged to express ACTB-mNeon or ACTB-mStayGold were incubated with 100 nM SiR-DNA for 2 h prior to imaging using a Nikon microscope outfitted with a CFI Plan Apo IR 60XC WI water immersion objective (NA 1.27), CrestOptics-X-Light-V2 spinning-disk microscope. For each field of view, samples were imaged every min for 20 min at 0.5 μm Z-step size for 15 μm total first using the 635 nm laser to image SiR-DNA followed by a 437 nm laser to image the fluorescently tagged proteins. The 437 nm laser was left on between time points to continuously illuminate the sample. Finally, volumetric live-cell super-resolution microscopy was performed on the Nikon CrestOptics-X-Light-DeepSIM microscope in SIM mode using the. To capture mStayGold-ACTB dynamics, 17 structured illumination images were captured at each z-step of 0.3 μm totaling 6 μm axially. Imaging occurred every 5 minutes for a total of 4 h. SIM reconstruction was then carried out to achieve super-resolution. Displayed are maximum intensity projections.

### Software & Analysis

General image processing was carried out with Nikon NISelements or ImageJ^58^ for deconvolution, maximum intensity projections, contrast modifications, and counting quantifications. For the photobleaching quantification, unprocessed, maximum intensity projections were analyzed using the following pipeline. Nuclear masks were obtained using the Stardist^59^ plug-in in ImageJ. Background was subtracted from the DNA channel using a 100-pixel rolling ball method, resized to 60% and analyzed using the default parameters in Stardist. Nuclear masks were imported into Cellprofiler^60^ with the ACTB-fusion channel images. Nuclei were filtered using a size cut off to exclude spurious objects and final nuclear objects were used as a seeding point to detect the cell boundaries of the ACTB-fusion channel using the Propagation method. The background of the ACTB-fusion channel was estimated using the lower quartile of the entire image and subtracted from the channel. The corrected ACTB-fusion channel was masked using the cell boundary objects and the total intensities were measured. The average object total intensity per image was plotted. Four fields of views were analyzed for each biological replicate (n = 3). General DNA sequence viewing, annotation, primer design, and cloning design was carried out in SnapGene 7.1.2. Flow cytometry analysis and visualization was performed in BD FlowJo 10.10. Samples were gated to exclude debris and cell doublets and +GFP fluorescence gates were determined from negative, unedited lines as well as auto fluorescent controls. Data analysis and visualization was performed in GraphPad Prism 10.2. Cellular models and diagrams were made in part by BioRender and further modified in Adobe Illustrator.

## Abbreviations

CRISPR: clustered regularly interspaced short palindromic repeats
sgRNA: single guide RNA
PAM: protospacer adjacent motif
NHEJ: non-homologous end joining
HDR: homology directed repair
MMEJ: microhomology-mediated end joining
DSB: double-strand break
Indels: insertion-deletion
GOI: gene of interest
bp: base pair
kb: kilobase
RNP: ribonucleoprotein

## Acknowledgements

We thank members of the Pelletier Lab for their scientific feedback during the project.

R.P. was partially funded by the Lunenfeld Tanenbaum Research Institute Studentships at Sinai Health System. A.S. is partially funded by the Hold’em For Life Oncology Fellowship. A.S. and A.C.E are partially funded by a CIHR Post Doctoral Grant (#187836 and #181763, respectively). H.D.M.W is funded by a CIHR Project Grant (#156297). H.D.M.W is a Tier II Canada Research Chair in Mechanisms of Genome Stability (#950-231487). The remainder of this work was funded by CIHR Foundation (FDN # 167279) and Krembil Foundation grants to L.P. L.P. is a Tier 1 Canada Research Chair in Centrosome Biogenesis and Function. This work was supported by the Lunenfeld-Tanenbaum Research Institute Flow Cytometry Core and The Network Biology Collaborative Centre which are funded through Canada Foundation for Innovation, the Ontario Government, and Genome Canada and Ontario Genomics (OGI-139) and the Nikon Center of Excellence at the Lunenfeld-Tanenbaum Research Institute. The schematics and models found in this manuscript were created in part by using BioRender.

## Author contributions

R.P., A.S., and J.M.T. developed the reagents and protocol. R.P., A.S., and J.M.T. designed the experiments. R.P., A.S., L.M., A.C.E., W-H.H., and J.M.T. performed the experiments. H.D.M.W purified and provided the Cas9 protein. L.P supervised the project. R.P. and L.P. wrote the manuscript with feedback from A.S., L.M., A.C.E., W-H.H., J.M.T., and H.D.M.W.

## Competing Interests

The authors declare no competing interests.

---

**Supplemental Figure 1.**
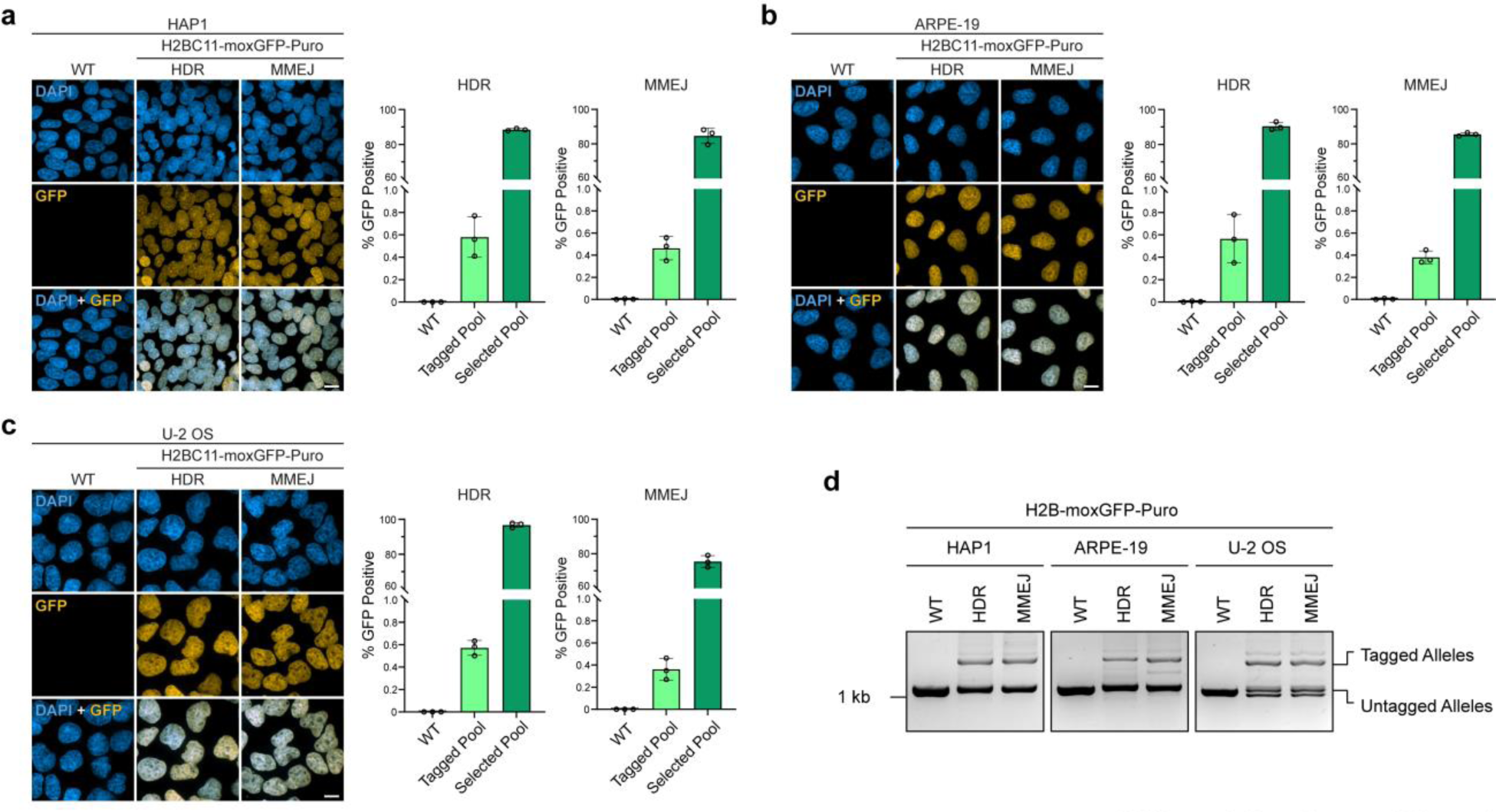
Editing, enrichment, and validation of fluorescent knock-ins with qTAG cassettes across different human cell lines. (a, b, c) Representative images of WT, HDR-integrated, and MMEJ-integrated H2BC11-moxGFP-Puro tagged HAP1, ARPE-19, and U-2 OS cells co-stained with DAPI (Left). Flow cytometry quantifications of GFP-positive cells, based on three distinct biological replicates with each measurement encompassing 200,000 cells (Right). (d) Genomic PCR outside the homology arms probing for locus-specific integration of the qTAG cassette in HAP1, ARPE-19, and U-2 OS cells. Scale bars: 10 µm.

**Supplemental Figure 2.**
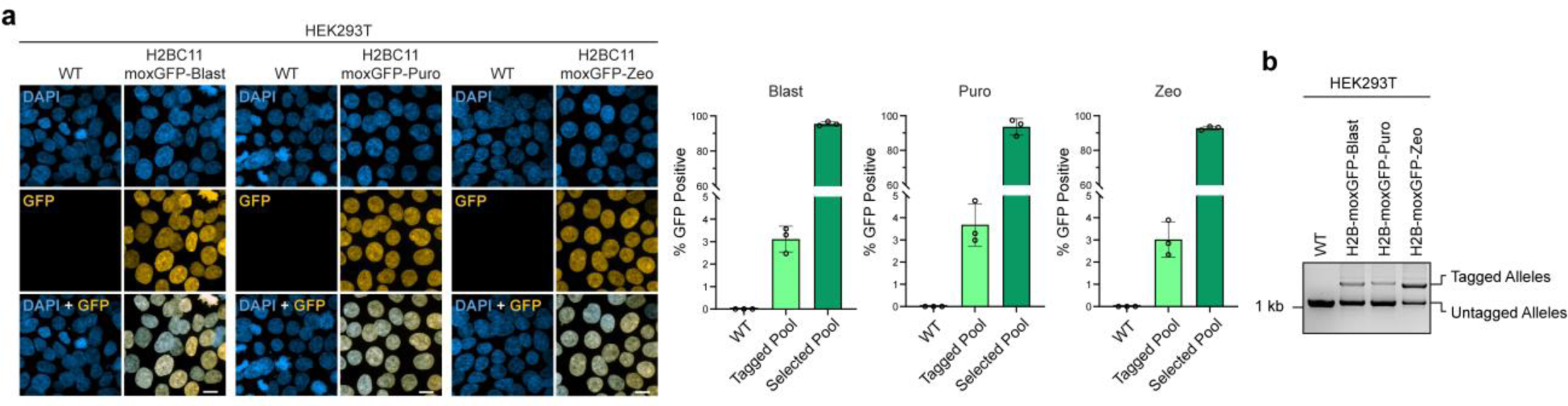
Editing, enrichment, and validation of fluorescent knock-ins with qTAG cassettes containing various selection markers. (a) Representative images of WT, selected H2BC11-moxGFP-Blast, selected H2BC11-moxGFP-Puro, and selected H2BC11-moxGFP-Zeo HEK293T cells co-stained with DAPI (Left). Flow cytometry quantifications of GFP-positive cells, based on three distinct biological replicates with each measurement encompassing 200,000 cells (Right). (b) Genomic PCR outside the homology arms probing for locus-specific integration of the qTAG cassettes with alternative mammalian selectable markers in HEK293T cells. Scale bars: 10 µm.

**Supplemental Figure 3.**
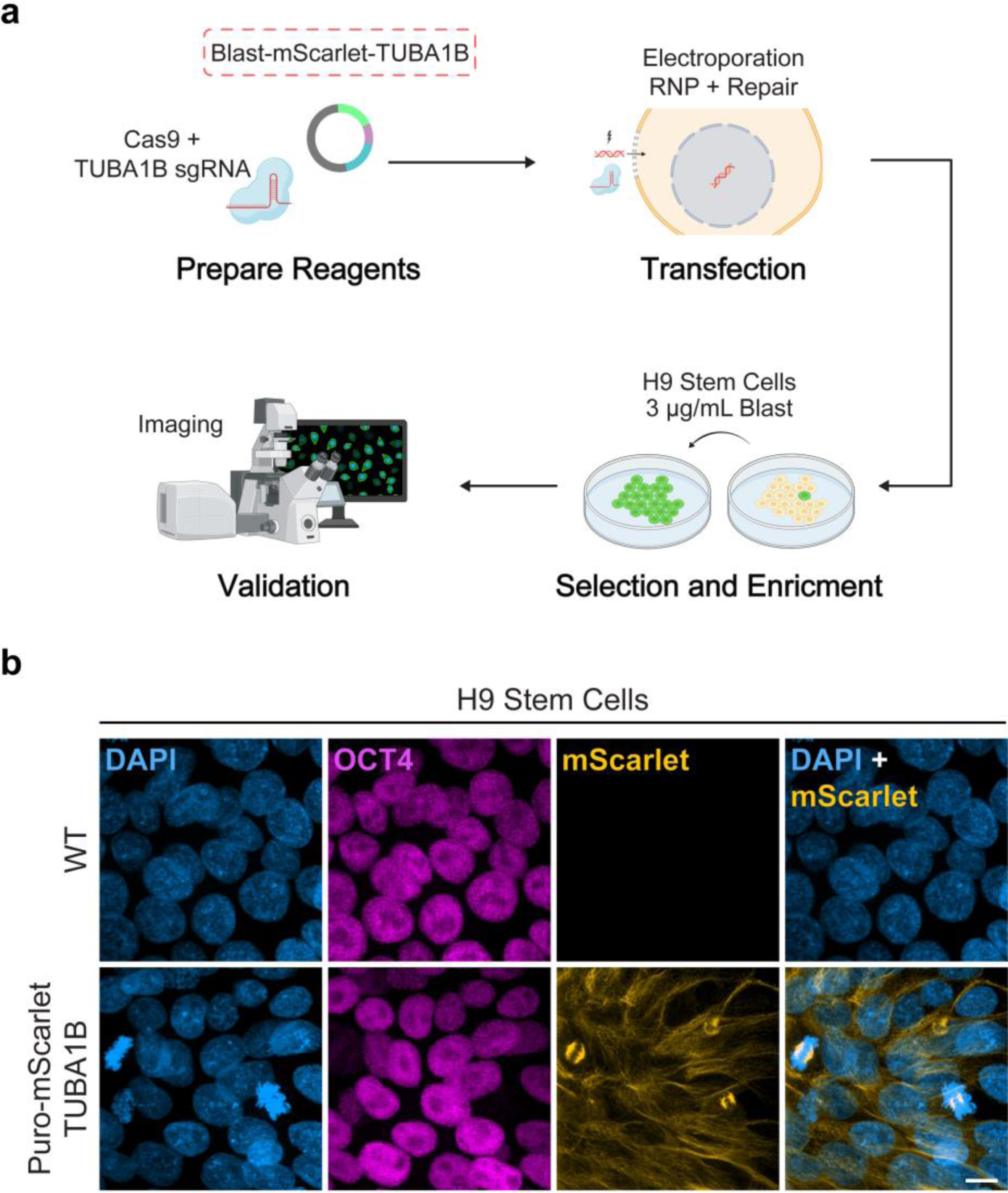
Fluorescent knock-in of H9 human embryonic stem cells with a qTAG cassette. (g) Overview of an alternate strategy using electroporation and RNPs to edit the tubulin TUBA1B gene with a qTAG-Blast-mScarlet cassette in H9 stem cells. (h) Representative images of WT and Blast-mScarlet-TUBA1B H9 cells co-stained with DAPI and probed for pluripotency marker OCT4. Scale bars: 10 µm.

Supplemental Data 1. A collection of files highlighting the design, cloning sequences, primer locations, and final plasmid products using qTAG cassettes targeting N-terminus or C-terminus insertions. (a) The provided sequence files are fully annotated and highlight the design of a C-terminal insertion of a qTAG-moxGFP-Puro cassette into the H2BC11 gene. They include a genomic loci design file, HDR and MMEJ homology arm designs, along with the final cloned CRISPR and repair plasmids. (b) The provided sequence files are fully annotated and highlight the design of a N-terminal insertion of a qTAG-Puro-Neon cassette into the ACTB gene. All of the corresponding design and final plasmid files are also included for this example.

## Notes

### Competing Interest Statement

The authors have declared no competing interest.

### Summary of Updates

This manuscript has been changed from a protocol format to a study format. This includes changes to the data as well as writing structure.

## References

1 Farhan, H., Weiss, M., Tani, K., Kaufman, R. J. & Hauri, H. P. Adaptation of endoplasmic reticulum exit sites to acute and chronic increases in cargo load. EMBO J 27, 2043–2054 (2008). 10.1038/emboj.2008.136

2 Vavouri, T., Semple, J. I., Garcia-Verdugo, R. & Lehner, B. Intrinsic protein disorder and interaction promiscuity are widely associated with dosage sensitivity. Cell 138, 198–208 (2009). 10.1016/j.cell.2009.04.029

3 Makanae, K., Kintaka, R., Makino, T., Kitano, H. & Moriya, H. Identification of dosage-sensitive genes in Saccharomyces cerevisiae using the genetic tug-of-war method. Genome Res 23, 300–311 (2013). 10.1101/gr.146662.112

4 Tang, Y. C. & Amon, A. Gene copy-number alterations: a cost-benefit analysis. Cell 152, 394–405 (2013). 10.1016/j.cell.2012.11.043

5 Bolognesi, B. et al. A Concentration-Dependent Liquid Phase Separation Can Cause Toxicity upon Increased Protein Expression. Cell Rep 16, 222–231 (2016). 10.1016/j.celrep.2016.05.076

6 Jinek, M. et al. A programmable dual-RNA-guided DNA endonuclease in adaptive bacterial immunity. Science 337, 816–821 (2012). 10.1126/science.1225829

7 Paques, F. & Haber, J. E. Multiple pathways of recombination induced by double-strand breaks in Saccharomyces cerevisiae. Microbiol Mol Biol Rev 63, 349–404 (1999). 10.1128/MMBR.63.2.349-404.1999

8 Urnov, F. D., Rebar, E. J., Holmes, M. C., Zhang, H. S. & Gregory, P. D. Genome editing with engineered zinc finger nucleases. Nat Rev Genet 11, 636–646 (2010). 10.1038/nrg2842

9 Moore, J. K. & Haber, J. E. Cell cycle and genetic requirements of two pathways of nonhomologous end-joining repair of double-strand breaks in Saccharomyces cerevisiae. Mol Cell Biol 16, 2164–2173 (1996). 10.1128/MCB.16.5.2164

10 Bibikova, M., Golic, M., Golic, K. G. & Carroll, D. Targeted chromosomal cleavage and mutagenesis in Drosophila using zinc-finger nucleases. Genetics 161, 1169–1175 (2002). 10.1093/genetics/161.3.1169

11 Chang, H. H. Y., Pannunzio, N. R., Adachi, N. & Lieber, M. R. Non-homologous DNA end joining and alternative pathways to double-strand break repair. Nat Rev Mol Cell Biol 18, 495–506 (2017). 10.1038/nrm.2017.48

12 Hart, T. et al. High-Resolution CRISPR Screens Reveal Fitness Genes and Genotype-Specific Cancer Liabilities. Cell 163, 1515–1526 (2015). 10.1016/j.cell.2015.11.015

13 Jasin, M. & Rothstein, R. Repair of strand breaks by homologous recombination. Cold Spring Harb Perspect Biol 5, a012740 (2013). 10.1101/cshperspect.a012740

14 Sakuma, T., Nakade, S., Sakane, Y., Suzuki, K. T. & Yamamoto, T. MMEJ-assisted gene knock-in using TALENs and CRISPR-Cas9 with the PITCh systems. Nat Protoc 11, 118–133 (2016). 10.1038/nprot.2015.140

15 Sfeir, A. & Symington, L. S. Microhomology-Mediated End Joining: A Back-up Survival Mechanism or Dedicated Pathway? Trends Biochem Sci 40, 701–714 (2015). 10.1016/j.tibs.2015.08.006

16 Taleei, R. & Nikjoo, H. Biochemical DSB-repair model for mammalian cells in G1 and early S phases of the cell cycle. Mutat Res 756, 206–212 (2013). 10.1016/j.mrgentox.2013.06.004

17 Fueller, J. et al. CRISPR-Cas12a-assisted PCR tagging of mammalian genes. J Cell Biol 219 (2020). 10.1083/jcb.201910210

18 Artegiani, B. et al. Fast and efficient generation of knock-in human organoids using homology-independent CRISPR-Cas9 precision genome editing. Nat Cell Biol 22, 321–331 (2020). 10.1038/s41556-020-0472-5

19 Roberts, B. et al. Systematic gene tagging using CRISPR/Cas9 in human stem cells to illuminate cell organization. Mol Biol Cell 28, 2854–2874 (2017). 10.1091/mbc.E17-03-0209

20 He, X. et al. Knock-in of large reporter genes in human cells via CRISPR/Cas9-induced homology-dependent and independent DNA repair. Nucleic Acids Res 44, e85 (2016). 10.1093/nar/gkw064

21 Perez-Leal, O. et al. Multiplex Gene Tagging with CRISPR-Cas9 for Live-Cell Microscopy and Application to Study the Role of SARS-CoV-2 Proteins in Autophagy, Mitochondrial Dynamics, and Cell Growth. CRISPR J 4, 854–871 (2021). 10.1089/crispr.2021.0041

22 Supharattanasitthi, W., Carlsson, E., Sharif, U. & Paraoan, L. CRISPR/Cas9-mediated one step bi-allelic change of genomic DNA in iPSCs and human RPE cells in vitro with dual antibiotic selection. Sci Rep 9, 174 (2019). 10.1038/s41598-018-36740-2

23 Lin, D. W. et al. Microhomology-based CRISPR tagging tools for protein tracking, purification, and depletion. J Biol Chem 294, 10877–10885 (2019). 10.1074/jbc.RA119.008422

24 Nabet, B. et al. The dTAG system for immediate and target-specific protein degradation. Nat Chem Biol 14, 431–441 (2018). 10.1038/s41589-018-0021-8

25 Nakamae, K. et al. Establishment of expanded and streamlined pipeline of PITCh knock-in - a web-based design tool for MMEJ-mediated gene knock-in, PITCh designer, and the variations of PITCh, PITCh-TG and PITCh-KIKO. Bioengineered 8, 302–308 (2017). 10.1080/21655979.2017.1313645

26 Pelletier, J. & Sonenberg, N. Internal initiation of translation of eukaryotic mRNA directed by a sequence derived from poliovirus RNA. Nature 334, 320–325 (1988). 10.1038/334320a0

27 Jang, S. K. et al. A segment of the 5’ nontranslated region of encephalomyocarditis virus RNA directs internal entry of ribosomes during in vitro translation. J Virol 62, 2636–2643 (1988). 10.1128/JVI.62.8.2636-2643.1988

28 Kim, J. H. et al. High cleavage efficiency of a 2A peptide derived from porcine teschovirus-1 in human cell lines, zebrafish and mice. PLoS One 6, e18556 (2011). 10.1371/journal.pone.0018556

29 Luke, G. A. et al. Occurrence, function and evolutionary origins of ‘2A-like’ sequences in virus genomes. J Gen Virol 89, 1036–1042 (2008). 10.1099/vir.0.83428-0

30 Sauer, B. & Henderson, N. Site-specific DNA recombination in mammalian cells by the Cre recombinase of bacteriophage P1. Proc Natl Acad Sci U S A 85, 5166–5170 (1988). 10.1073/pnas.85.14.5166

31 Albert, H., Dale, E. C., Lee, E. & Ow, D. W. Site-specific integration of DNA into wild-type and mutant lox sites placed in the plant genome. Plant J 7, 649–659 (1995). 10.1046/j.1365-313x.1995.7040649.x

32 Zhang, Z. & Lutz, B. Cre recombinase-mediated inversion using lox66 and lox71: method to introduce conditional point mutations into the CREB-binding protein. Nucleic Acids Res 30, e90 (2002). 10.1093/nar/gnf089

33 Ando, R. et al. StayGold variants for molecular fusion and membrane-targeting applications. Nat Methods (2023). 10.1038/s41592-023-02085-6

34 Shaner, N. C. et al. A bright monomeric green fluorescent protein derived from Branchiostoma lanceolatum. Nat Methods 10, 407–409 (2013). 10.1038/nmeth.2413

35 Costantini, L. M. et al. A palette of fluorescent proteins optimized for diverse cellular environments. Nat Commun 6, 7670 (2015). 10.1038/ncomms8670

36 Mo, G. C. H., Posner, C., Rodriguez, E. A., Sun, T. & Zhang, J. A rationally enhanced red fluorescent protein expands the utility of FRET biosensors. Nat Commun 11, 1848 (2020). 10.1038/s41467-020-15687-x

37 Bindels, D. S. et al. mScarlet: a bright monomeric red fluorescent protein for cellular imaging. Nat Methods 14, 53–56 (2017). 10.1038/nmeth.4074

38 Oliinyk, O. S. et al. Single-domain near-infrared protein provides a scaffold for antigen-dependent fluorescent nanobodies. Nat Methods 19, 740–750 (2022). 10.1038/s41592-022-01467-6

39 Kubitz, L. et al. Engineering of ultraID, a compact and hyperactive enzyme for proximity-dependent biotinylation in living cells. Commun Biol 5, 657 (2022). 10.1038/s42003-022-03604-5

40 Branon, T. C. et al. Efficient proximity labeling in living cells and organisms with TurboID. Nat Biotechnol 36, 880–887 (2018). 10.1038/nbt.4201

41 Hirano, M. et al. A highly photostable and bright green fluorescent protein. Nat Biotechnol 40, 1132–1142 (2022). 10.1038/s41587-022-01278-2

42 Ivorra-Molla, E. et al. A monomeric StayGold fluorescent protein. Nat Biotechnol (2023). 10.1038/s41587-023-02018-w

43 Zhang, H. et al. Bright and stable monomeric green fluorescent protein derived from StayGold. Nat Methods (2024). 10.1038/s41592-024-02203-y

44 Guderian, G., Westendorf, J., Uldschmid, A. & Nigg, E. A. Plk4 trans-autophosphorylation regulates centriole number by controlling betaTrCP-mediated degradation. J Cell Sci 123, 2163–2169 (2010). 10.1242/jcs.068502

45 Coelho, P. A. et al. Over-expression of Plk4 induces centrosome amplification, loss of primary cilia and associated tissue hyperplasia in the mouse. Open Biol 5, 150209 (2015). 10.1098/rsob.150209

46 Kojima, K. et al. Mdm2 inhibitor Nutlin-3a induces p53-mediated apoptosis by transcription-dependent and transcription-independent mechanisms and may overcome Atm-mediated resistance to fludarabine in chronic lymphocytic leukemia. Blood 108, 993–1000 (2006). 10.1182/blood-2005-12-5148

47 Liang, X., Potter, J., Kumar, S., Ravinder, N. & Chesnut, J. D. Enhanced CRISPR/Cas9-mediated precise genome editing by improved design and delivery of gRNA, Cas9 nuclease, and donor DNA. J Biotechnol 241, 136–146 (2017). 10.1016/j.jbiotec.2016.11.011

48 Zhong, H. et al. High-fidelity, efficient, and reversible labeling of endogenous proteins using CRISPR-based designer exon insertion. Elife 10 (2021). 10.7554/eLife.64911

49 Cho, N. H. et al. OpenCell: Endogenous tagging for the cartography of human cellular organization. Science 375, eabi6983 (2022). 10.1126/science.abi6983

50. Willems, J., et al. ORANGE: A CRISPR/Cas9-based genome editing toolbox for epitope tagging of endogenous proteins in neurons. PLoS Biol 18, e3000665 (2020). 10.1371/journal.pbio.3000665

51 Liu, Z. et al. Systematic comparison of 2A peptides for cloning multi-genes in a polycistronic vector. Sci Rep 7, 2193 (2017). 10.1038/s41598-017-02460-2

52 Szczesny, R. J. et al. Versatile approach for functional analysis of human proteins and efficient stable cell line generation using FLP-mediated recombination system. PLoS One 13, e0194887 (2018). 10.1371/journal.pone.0194887

53 Labun, K. et al. CHOPCHOP v3: expanding the CRISPR web toolbox beyond genome editing. Nucleic Acids Res 47, W171–W174 (2019). 10.1093/nar/gkz365

54 Cong, L. et al. Multiplex genome engineering using CRISPR/Cas systems. Science 339, 819–823 (2013). 10.1126/science.1231143

55 Zuris, J. A. et al. Cationic lipid-mediated delivery of proteins enables efficient protein-based genome editing in vitro and in vivo. Nat Biotechnol 33, 73–80 (2015). 10.1038/nbt.3081

56 Anders, C. & Jinek, M. In vitro enzymology of Cas9. Methods Enzymol 546, 1–20 (2014). 10.1016/B978-0-12-801185-0.00001-5

57 Hu, Z. et al. Customized one-step preparation of sgRNA transcription templates via overlapping PCR Using short primers and its application in vitro and in vivo gene editing. Cell Biosci 9, 87 (2019). 10.1186/s13578-019-0350-7

58. Schindelin, J., et al. Fiji: an open-source platform for biological-image analysis. Nat Methods 9, 676-682 (2012). 10.1038/nmeth.2019

59 Schmidt, U., Weigert, M., Broaddus, C. & Myers, G. in *Medical Image Computing and Computer Assisted Intervention – MICCAI 2018 Lecture Notes in Computer Science* Ch. Chapter 30, 265–273 (2018).

60 Stirling, D. R. et al. CellProfiler 4: improvements in speed, utility and usability. BMC Bioinformatics 22, 433 (2021). 10.1186/s12859-021-04344-9

